# Patient-Specific 3D Heart-On-a-Chip Model of Dilated Cardiomyopathy with Embedded Bead-Based Mapping of Tissue Contractility

**DOI:** 10.64898/2025.12.20.695633

**Authors:** Ali Mousavi, Ludovic Mouttet, Shihao Cui, Yasaman Hekmatnia, Mehran Mottahedi, Ida Derish, Naimeh Rafatian, Mark Aurousseau, Gregor Andelfinger, Renzo Cecere, Houman Savoji

## Abstract

Dilated cardiomyopathy (DCM) is the leading cause of heart transplantation, with a 50% risk of progression to heart failure within five years. Conventional disease modeling approaches fail to recapitulate the sophisticated function of the human heart. Alternatively, heart-on-a-chip (HOC) platforms enable real-time monitoring of disease progression and drug responses using miniaturized engineered heart tissues. Here, we developed a functional HOC model using patient-specific human induced pluripotent stem cells (hiPSCs), reprogrammed from the patients’ blood samples. The chip contains two cell-seeding chambers with flexible silicone pillars to support tissue formation. Healthy and DCM hiPSCs were differentiated into cardiomyocytes, combined with an optimized ratio of human cardiac fibroblasts, encapsulated in a fibrin/Geltrex hydrogel (containing fluorescent beads), and seeded in the device chambers. The tissue gradually compacted and started beating spontaneously. Immunofluorescence assay revealed structural abnormalities in DCM tissues, including reduced cell alignment and elongation. The tissue functional responses (e.g., calcium transients and beating) were investigated after 2 weeks of culture, revealing ventricular tachycardia in DCM tissue and highlighting functional hallmarks of the disease. Finally, the model was validated using a drug with known inotropic and chronotropic effects (i.e., norepinephrine). Our platform demonstrated great potential in drug screening, disease modeling, and personalized medicine.

## 1. Introduction

Cardiovascular disease (CVD) remains the primary cause of mortality worldwide, accounting for approximately 17.9 million deaths annually, according to the World Health Organization [1]. Among CVDs, dilated cardiomyopathy (DCM) is a non-ischemic cardiac disorder showing structural and functional abnormalities in the myocardium, and it is generally characterized by left or biventricular dilation and systolic impairment [2]. DCM is known to be the most common type of cardiomyopathy, the leading cause of heart failure, and the primary indication for heart transplantation [3]. It is categorized as primary (e.g., idiopathic, familial) or secondary, with the latter potentially arising from various external factors, including hypertension, ischemic heart disease, coronary artery disease, metabolic dysfunctions, and drug-induced disorders (e.g., doxorubicin) [4]. Despite the diverse etiology, idiopathic DCM is the most prevalent form, making up 60-70% of cases, in which no definitive clinical cause or genetic origin can be identified. Therefore, a comprehensive assessment is often required to accurately determine the underlying etiology [4–6].

One significant challenge in cardiovascular disease modeling is the lack of a non-invasive method to isolate human cardiomyocytes (CMs) from patients or grow them *in vitro* due to their limited proliferative capacity [7]. Since the breakthrough discovery in 2007, human induced pluripotent stem cells (hiPSCs) have revolutionized this field and have become a promising alternative for creating patient-specific disease models *in vitro* [8]. These stem cells could be derived from somatic cells (e.g., skin fibroblasts or peripheral blood) and offer several advantages, such as ease of expansion and differentiation into various cell lines (e.g., CMs), providing an unlimited source of patient-specific CMs *in vitro* [9]. hiPSC-derived CMs (hiPSC-CMs) possess the specific features of human CMs and maintain the donor’s genetic background and the disease hallmarks (e.g., arrhythmia), making them an ideal candidate for cardiovascular disease modeling [10, 11].

Traditional disease modeling methods (e.g., animal models and two-dimensional (2D) monolayer cell models) lack physiological relevance to the human heart due to interspecies genetic differences (e.g., contraction rate, ion channel dynamics, and heart size) [12]. As a versatile alternative, heart-on-a-chip (HOC) has been introduced to mimic the intricate three-dimensional (3D) architecture and function of the heart, which could be utilized for preclinical drug screening [13]. One of the major advantages of HOC models is the ability to continuously measure tissue-scale contractility. Among conventional techniques, traction force microscopy (TFM) has been used to quantify the forces exerted by single cells or cell sheets on a substrate (e.g., polydimethylsiloxane (PDMS), polyacrylamide gels) by tracking the movement of beads embedded within it [14]. This method has primarily been applied in 2D, which restricts its use for measuring contractility in 3D tissues. One of the most common 3D HOC models is engineered heart tissue (EHT), formed through compaction between two defined anchoring structures, such as pillars and wires [15, 16]. In these systems, the tissue contractile activity could be measured based on the deflection of the pillars during contraction [17]. However, this method does not capture force heterogeneity across the tissue and only reflects the overall tissue contractility near the pillar.

To overcome this limitation, we have developed a 3D TFM model by embedding fluorescent beads within 3D EHTs. The chip features pillar pairs in each chamber, enabling measurement of contractile forces via pillar deflection. Unlike conventional 2D TFM models, in which beads are embedded in a secondary substrate, the fluorescent beads were directly incorporated into the cell-hydrogel mixture prior to tissue seeding and formation. This method enabled real-time mapping of tissue contractile activity throughout the tissue, complementing force calculations based on pillar deflection. The hiPSC-CMs, derived from idiopathic DCM patients and healthy controls, were used in this platform, and EHTs from both groups were functionally characterized based on contractile properties, calcium dynamics, IF staining of cardiac-specific biomarkers, and response to pharmacological interventions. Disease modeling was achieved by comparing the functional performance of DCM and control EHTs (**Fig. 1**). To the best of our knowledge, this is the first HOC model of DCM that combines embedded bead-based 3D TFM and pillar deflection methods to map contractility and assess arrhythmia in patient-specific tissues.

**Fig. 1.**
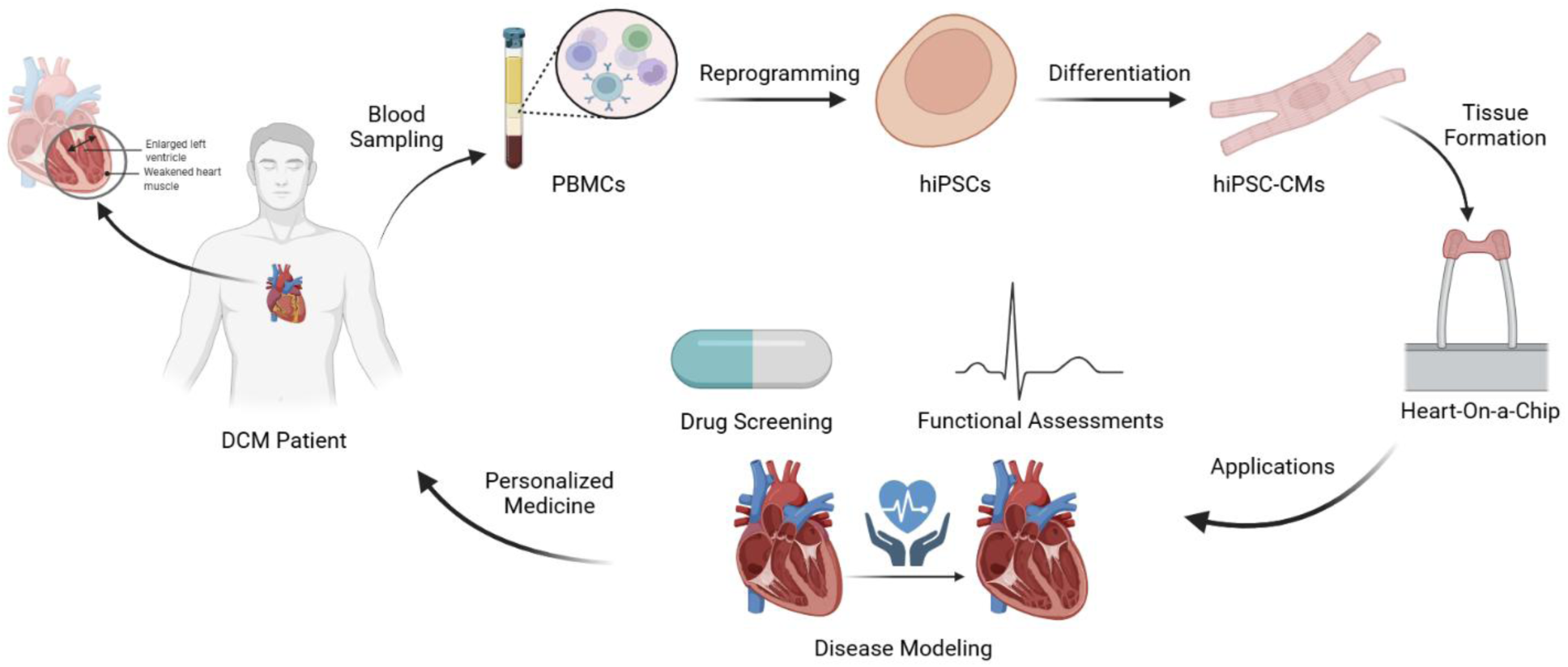
Schematic overview of the study outlining the generation of patient-specific hiPSCs from blood samples, differentiation into CMs, and formation of 3D cardiac tissues within an HOC platform. The workflow includes functional assessments (e.g., contractile force measurements and calcium transient analysis), followed by validation of the system for drug screening and disease modeling toward personalized medicine applications (generated with BioRender).

## 2. Materials and Methods

### 2.1. Materials

Dulbecco’s modified Eagle medium (DMEM)/F12 culture medium (#11320033), Dulbecco’s phosphate buffered saline (DPBS; without calcium and magnesium), RPMI 1640 medium (#11875119), glucose-free RPMI 1640 medium (#11879020), B-27 supplement (#17504001), B-27 supplement minus insulin (#A1895602), KnockOut Serum Replacement (KOSR, #10828028), fetal bovine serum (FBS, #16140089), penicillin/streptomycin (Pen Strep, #15140122), Geltrex basement membrane matrix (#A1413302), GlutaMAX (#35050061) were purchased from Gibco, Canada. Claycomb medium (#51800C), fibrinogen (#F8630), thrombin (#T4648), aprotinin (#A3428), sodium L-lactate (#L7022), monoclonal anti-α-actinin (sarcomeric) antibody (#A7811), Pluronic-F127, bovine serum albumin (BSA), neutral buffered formalin (NBF, 10%), and norepinephrine (NE) were supplied by Sigma Aldrich, Canada. Fluo-4 NW calcium assay kit (#F36206), Fluoromount-G mounting medium (#00-4958-02), 4′,6-aiamidino-2-phenylindole dihydrochloride (DAPI, #D1306), ActinRed 555 ReadyProbes reagent (#R37112), NucBlue Live ReadyProbe Reagent (Hoechst 33342, #R37605), cardiac Troponin T (cTnT) monoclonal antibody (#MA5-12960), sarcoplasmic/endoplasmic reticulum calcium ATPase (SERCA2a) monoclonal antibody (#MA3-919), connexin 43 (Cx43) polyclonal antibody (#71-0700), GATA4 polyclonal antibody (#PA1-102), and secondary antibodies (including goat anti-mouse Alexa Fluor 488 (#A11001), donkey anti-rabbit Alexa Fluor 488 (#A21206), and goat anti-mouse Alexa Fluor 647 (#A21235)) were purchased from Invitrogen, USA. Phalloidin-iFluor 488 reagent (#ab176753) and anti-Myosin Light Chain 2 (MLC-2v) antibody (#ab79935) were obtained from Abcam, UK. Nuclear pluripotency markers octamer-binding transcription factor 4 (OCT4) and homeobox protein NANOG, and surface markers stage-specific embryonic antigen-4 (SSEA-4) and tumor-related antigen-1-60 (TRA-1-60) were supplied by Cell Signaling, USA. CHIR99021 (#S1263), IWP-2 (#S7085), and Rho kinase (ROCK) inhibitor (Y-27632 dihydrochloride, #S1049) were purchased from Selleck Chemicals, USA. Accutase (#07920) and mTeSR Plus medium (#100-0276) were obtained from STEMCELL Technologies, Canada.

### 2.2. Chip Design

The microphysiological systems, commercially branded as OMEGA^MP^, were provided by eNUVIO Inc. (Montréal, QC, Canada). These PDMS-based chips contain two cell-seeding chambers, each equipped with vertical millipillar pairs to promote tissue attachment and secure anchorage. These devices were specifically engineered to fit firmly within standard 12-well culture plates, ensuring seamless compatibility with existing lab protocols. The elliptical seeding chamber (8 mm × 4 mm × 2 mm) offers a surface area of 0.25 cm² and a maximum volume of 40 μl. The 3 mm-tall pillars have a tilted structure to prevent tissue detachment and enhance stability. Their diameter gradually decreases as they extend from the PDMS base, averaging 1 mm. The PDMS base beneath the chambers is 0.2 mm thick, providing a transparent layer that enables high-resolution live imaging at the microscopic level on the chip (**Fig. S1**).

### 2.3. Patient Selection and hiPSC Generation

Control and patient individuals were recruited from the cardiology clinic at the McGill University Health Centre. The control group consisted of either healthy relatives of patients or individuals without any known cardiopulmonary conditions. All participants provided informed consent to take part in the “Heart-in-a-Dish” project, which received approval from the Research Ethics Board (REB) under protocol HID-B/2020-6362. Each individual completed a questionnaire gathering details on biological sex, ethnicity, age, family history of cardiovascular disease, smoking habits, cholesterol levels, history of cancer diagnoses, current medications, and responses to a Genesis Praxy gender questionnaire. In this study, the control subject and DCM patient were selected based on matched background data, and both were middle-aged (40 to 45 years old) white French-Canadian females. The patient had idiopathic DCM with no additional clinical diagnoses. The patient-specific hiPSCs were generated by reprogramming their peripheral blood mononuclear cells (PBMCs) through the introduction of Yamanaka factors (i.e., Oct4, Sox2, Klf4, and L-Myc) using the Epi5^TM^ Episomal iPSC Reprogramming Kit (Thermo Fisher Scientific, USA) and the Neon Transfection System (Invitrogen, Canada), according to our previous report [18]. hiPSC colonies were characterized by immunofluorescence (IF) for nuclear pluripotency markers OCT4 and NANOG, as well as surface markers SSEA-4 and TRA-1-60. Furthermore, the functional ability of iPSCs to differentiate into the three germ layers (i.e., ectoderm, mesoderm, and endoderm) was assessed using a trilineage differentiation assay (#SC027B, R&D Systems, USA). The expression of orthodenticle homeobox 2 (OTX2), Brachyury, and sex determining region Y-box 17 (SOX17) was evaluated to confirm ectoderm, mesoderm, and endoderm differentiation, respectively.

### 2.4. hiPSC Differentiation to Cardiomyocytes

Patient-specific hiPSCs were differentiated into CMs using a modified GiWi protocol [19]. To establish a confluent monolayer, hiPSCs were seeded on Matrigel-coated 12-well plates, cultured in mTeSR Plus medium supplemented with 10 μM ROCK inhibitor, and incubated at 37°C with 5% CO₂. After 24 hours, the medium was replaced with fresh mTeSR plus, and cells were cultured for an additional 48 hours before initiating differentiation. On day 0, Wnt signaling was activated by replacing the culture medium with RPMI 1640 containing 2% (v/v) B-27 minus insulin and 12 μM CHIR99021, followed by a media change exactly 24 hours later. On day 1, fresh RPMI/B27 minus insulin was added, and on day 3, 5 μM IWP-2 was introduced to inhibit Wnt signaling. Medium changes with fresh RPMI/B27 minus insulin were performed on days 5 and 7. From day 9 onward, cells were cultured in RPMI 1640 supplemented with 2% (v/v) B-27, with an additional media change on day 11. CMs typically started beating spontaneously between days 7 and 10 and could be observed under bright-field microscopy. On day 13, metabolic selection was carried out by switching to glucose-free RPMI 1640 supplemented with 2% (v/v) B-27 and 4 mM sodium L-lactate for 4 days, with a media refresh after 2 days. After purification, hiPSC-CMs were maintained in RPMI/B27 media for an additional 24 h before cell dissociation and encapsulation.

### 2.5. Differentiation Characterizations

#### 2.5.1. Beating Assessments

The beating of hiPSC-CMs in control (CTRL) and patient (DCM) lines was recorded with an Olympus IX51 microscope (Olympus, Japan) on days 13 (before purification) and 18 (after purification, before cell dissociation, and before encapsulation). The videos were further quantified using a custom-built MATLAB script based on the particle image velocimetry (PIV) method, and the beating graphs were depicted as monolayer displacement versus time. Beating frequency, as beats per minute (BPM), and maximum displacements were quantified based on these graphs. Cell displacement was also mapped in both experimental groups, showing the relative changes in displacement across different parts of the culture plates.

#### 2.5.2. Immunocytochemistry

The CTRL and DCM iPSC-CMs were also characterized by immunocytochemistry (ICC) of cardiac-specific biomarkers, including cTnT, SERCA2a, Cx43, and GATA4. Briefly, the cells were fixed in 10% formalin solution for 15 min, followed by 3 washes with DPBS for 5 min each. The cells were further permeabilized in 0.5% Triton X-100 solution and then placed into a blocking buffer containing 3% BSA for 1 h. The primary antibody solutions were prepared by dilution (1:500 for Cx43, 1:150 for SERCA2a, 1:200 for GATA4, and 1:400 for cTnT) into the blocking buffer, and the cells were incubated in these solutions at 4 ℃ overnight. After 3 washes with DPBS, the cells were incubated with secondary antibody solutions (1:500 diluted in blocking buffer) for 1 h at room temperature. The counterstaining for filamentous actin (F-actin) and nuclei was then carried out with ActinRed and NucBlue for 30 min at room temperature by adding 2 drops of each per 1ml DPBS. The samples were further washed 3 times with DPBS and then transferred to a microscopic slide using a mounting medium. The samples were further visualized with an ApoTome.2 confocal fluorescent microscopes (Carl Zeiss, Germany).

#### 2.5.3. Gene Expression

The gene expression of CTRL and DCM iPSC-CMs was analyzed using the quantitative reverse transcription polymerase chain reaction (qRT-PCR) technique. Total RNA was extracted using the RNeasy Plus Kit (Qiagen, Germany) following the manufacturer’s protocol. Complementary DNA (cDNA) was further synthesized using the iScript Advanced cDNA Synthesis Kit (Bio-Rad, USA) in accordance with the supplier’s instructions. The reaction mixture was prepared with the PowerUp SYBR Green Master Mix (Applied Biosystems, USA), and qRT-PCR was performed using the primers listed in Table S1. Gene expression levels were normalized to the reference gene GAPDH.

### 2.6. Human Cardiac Fibroblast Culture

Adult human ventricular cardiac fibroblasts (HCFs) isolated from healthy explanted hearts and transplanted upon purchase were purchased from a commercial vendor (#C-12375, Lot #495Z001.7, 43-year-old Caucasian female donor, PromoCell, Germany). The cryopreserved cells were received at passage 2 (P2), thawed according to the supplier’s recommended protocol, and cultured using Fibroblast Growth Medium 3 (#C-23025, PromoCell, Germany) for up to 10 passages in T75 tissue culture flasks. In this study, these cells were used for the experiments between P4 and P6 at 70-90% confluency (Figure S1).

### 2.7. Human EHT Formation

On day 18 post-differentiation, hiPSC-CMs were enzymatically detached and dissociated using Accutase solution for 20 minutes at 37 °C. An equal volume of cell culture medium was further added to neutralize enzyme activity, and the cell suspension was gently pipetted to ensure complete dissociation into single cells. Furthermore, cells were centrifuged at 300 × g for 5 minutes, resuspended in the culture medium, and counted using a hemocytometer. hiPSC-CMs were then mixed with HCFs at a predetermined ratio. As a preliminary study, different ratios of hiPSC-CMs to HCFs (90:10, 80:20, and 70:30) were tested, and the optimal condition (80:20) was selected for further experiments. This cardiac cell mixture was then encapsulated in a fibrin/Geltrex matrix and seeded in the device chambers to form EHTs. Before cell encapsulation, all solutions were sterilized through 0.2 μm polyethersulfone (PES) syringe filters. Prior to cell seeding, the chips were carefully placed into each well of the 12-well plates using sterilized tweezers. The chips were then coated with 5% Pluronic F-127 solution in PBS and incubated overnight at 4°C to prevent unwanted cell attachment to the PDMS base. All following tissue formation procedures were performed on ice with pre-cooled pipette tips and hydrogel solutions.

The fibrinogen solution (4 mg/ml), 20% (v/v) Geltrex matrix, and fluorescent orange polystyrene beads (#2218, 3 μm nominal diameter, Phosphorex, USA) were added to the cardiac cell pellet to prepare the final cell-hydrogel suspension with a density of 40 million cells/ml. Prior to cell seeding, the antifouling coating agent was aspirated from the chips, and the device was washed once with cold PBS. Subsequently, the thrombin solution (at a concentration of 0.2 Units/mg of fibrinogen) was rapidly combined with the cell-hydrogel suspension, and 35 μL was evenly seeded into each chamber, ensuring no air bubbles were introduced. Following 15 minutes of incubation at 37 ℃ for completion of fibrin polymerization and tissue formation, the EHTs were cultured in RPMI medium supplemented with 2% (v/v) B-27, 1% Pen Strep, 10 μg/mL aprotinin, 10% (v/v) KOSR, and 10 μM ROCK inhibitor. The culture medium was replaced the following day with a fresh RPMI media (containing all supplements except KOSR and ROCK inhibitor) and subsequently refreshed every other day. Tissue formation, compaction, and contraction were regularly observed under an Olympus IX51 microscope (Olympus, Japan) for up to 2 weeks.

### 2.8. Immunofluorescence Staining

The EHTs were stained for cardiac structural and junctional markers (cTnT, Cx43, SERCA2a, MLC-2v, and α-actinin) [20]. The procedure followed the same steps as described in section 2.5.2. However, counterstaining for F-actin and Nuclei was performed using Phalloidin (1:1000) for 1 hour and DAPI solution (1:1000) for 20 minutes at room temperature, respectively. The EHTs were further imaged using the SP8-DLS Leica confocal microscope (Leica Biosystems, Germany). IF images were analyzed using ImageJ, and the density of each biomarker was calculated by measuring the fluorescence intensity values normalized to the corresponding DAPI intensity, as described elsewhere [21].

### 2.9. Contractility Measurements

The beating of EHTs was visualized on-chip using a spinning-disk confocal microscope (Carl Zeiss, Germany) with a 10x objective, based on bead movements and pillar deflections. The microscope chamber was equipped with a CO_2_ and temperature control system to maintain a stable environment for live tissue imaging at 37°C and 5% CO_2_. The videos were further analyzed using a custom MATLAB code described in Section 2.5.1, yielding bead movement graphs and maps. The contractile force was then calculated based on the force-displacement equation outlined in our previous report [22], and contractile stress was determined by normalizing the force to the cross-sectional area.

### 2.10. Calcium Imaging

Calcium handling was evaluated using the Fluo-4 NW calcium assay kit (Invitrogen, Canada) following the manufacturer’s recommended protocol. Briefly, EHTs were incubated with the prepared calcium indicator at 37°C for 45 minutes. The samples were then transferred to a high-speed spinning disk confocal microscope (Carl Zeiss, Germany), where fluorescent videos were recorded at 37°C and 5% CO₂ using a GFP filter cube. Calcium transient videos were analyzed using FIJI (NIH, USA), and calcium transient patterns were depicted by normalizing fluorescence intensity (F) to background fluorescence intensity (F_0_) over time, averaged across three defined regions of interest (ROIs).

### 2.11. Drug Testing

Norepinephrine (NE), a pharmaceutical compound with well-characterized cardiac effects, was used for drug testing [23]. The compound was diluted in culture medium, added to the EHTs at a final concentration of 100 μM, and incubated at 37°C for 45 minutes. Calcium handling and tissue contractility were measured before and after NE treatment following the protocols described in Sections 2.9 and 2.10.

### 2.12. Statistical Analysis

Statistical analyses were conducted using GraphPad Prism software version 9.0 (GraphPad Inc., USA). An unpaired t-test was used for this analysis since all comparisons involved two groups.

Data are presented as the mean ± S.D. with n = 6 (unless otherwise noted), and statistical significance was defined as a p-value of less than 0.05.

## 3. Results and Discussion

### 3.1. hiPSC Generation

In a personalized medicine approach to studying DCM, we have established a biorepository of patient-specific blood samples from individuals with DCM and healthy controls. Participants were recruited from diverse age groups, ethnic backgrounds, and sexualities across Canada. According to our biobank, developed under the “Heart-in-a-Dish” project, patients with DCM exhibit various etiologies, although the majority are classified as idiopathic. The patient-derived blood samples were reprogrammed into hiPSCs using episomal plasmids (**Fig. 2A**). These hiPSCs were validated by IF staining for pluripotency markers and confirmed through tri-lineage differentiation. As shown in **Fig. 2B**, both control and DCM-derived hiPSC lines expressed key pluripotency markers, including OCT4, NANOG, SSEA4, and TRA-1-60 (green), with nuclei counterstained in blue with Hoechst (**Fig. S3**). The tri-lineage differentiation assay demonstrated the potential of both hiPSC lines to differentiate into all three primary germ layers: ectoderm, mesoderm, and endoderm, which were depicted for specific markers OTX2, Brachyury, and SOX17, respectively, in green color (**Fig. 2C**). In these differentiated colonies, nuclei were stained in blue and F-actin in red, indicating cytoskeletal organization (**Fig. S4**). These results confirm the pluripotency and differentiation potential of both control and DCM iPSC lines, establishing a reliable foundation for subsequent cardiac differentiation and tissue modeling.

**Fig. 2.**
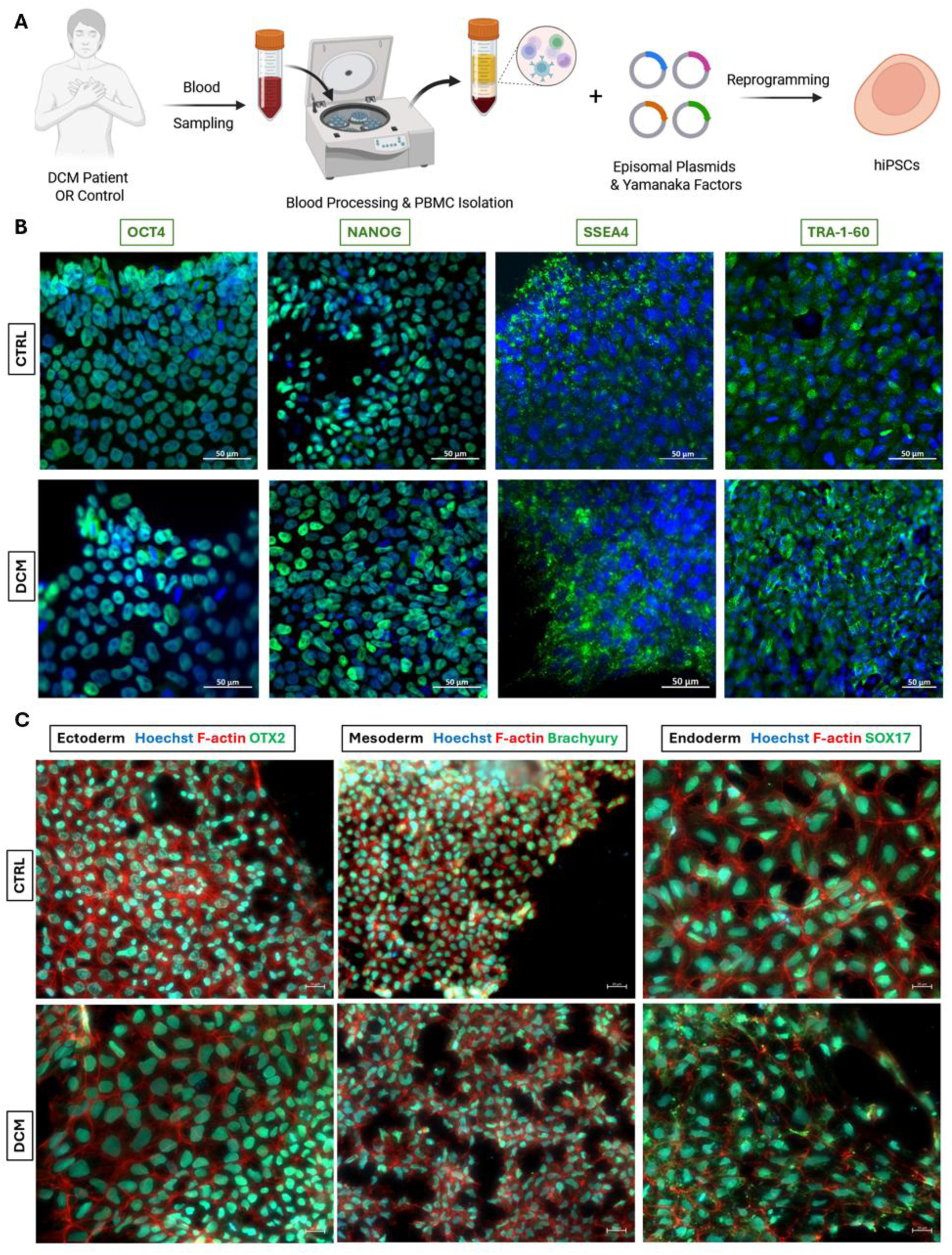
(A) Schematic representation of the reprogramming procedure to generate control- and patient-derived hiPSCs from blood samples via an episomal plasmid-based method. (B) IF staining confirmed the expression of key pluripotency markers, including OCT4 and NANOG (localized in nuclei), and SSEA4 and TRA-1-60 (surface markers), all shown in green. Nuclei were counterstained with Hoechst (blue). Scale bar = 50 μm. (C) Representative images of tri-lineage differentiation demonstrating the ability of hiPSC lines to generate three primary germ layers: ectoderm (OTX2), mesoderm (Brachyury), and endoderm (SOX17), each visualized in green. Nuclei and F-actin were depicted in blue and red, respectively. Scale bar = 20 μm.

### 3.2. Cardiac Differentiation

CM differentiation was achieved using a defined method based on the Wnt signaling pathway (**Fig. 3A**). Initially, CHIR99021, a GSK3β inhibitor, was added to activate the Wnt/β-catenin pathway, directing the differentiation towards mesodermal lineage. Afterward, IWP2 was applied as a Wnt pathway inhibitor, thereby facilitating cardiogenic differentiation. Following this biphasic modulation, spontaneously beating monolayers emerged in both control and DCM cell lines, demonstrating successful CM differentiation. The spontaneous beating of hiPSC-CMs usually began between days 7 and 9 of differentiation and was consistently maintained throughout the differentiation timeline. To enhance the CM purity, lactate-based metabolic selection was employed. This approach takes advantage of CMs’ distinct metabolic capacity compared to other cardiac cell types, as CMs can utilize lactate as an energy source, whereas most non-CMs cannot [24]. Consequently, the CMs were selectively enriched upon culturing the cells in a glucose-free, lactate-containing medium for 4 days. The spontaneous beating of hiPSC-CM monolayers was recorded on day 18 post-differentiation, immediately prior to cell dissociation and encapsulation (**Movie S1-2**). The videos were processed using a custom MATLAB code to generate displacement heatmaps (Fig. 3B). This approach enabled detection of spatial heterogeneity in contractile activity across the entire plate and quantitative analysis of displacement magnitudes. The DCM line exhibited significantly lower contractile strength (measured as monolayer displacement) and a higher beating frequency than the healthy control (**Fig. 3C-D**).

**Fig. 3.**
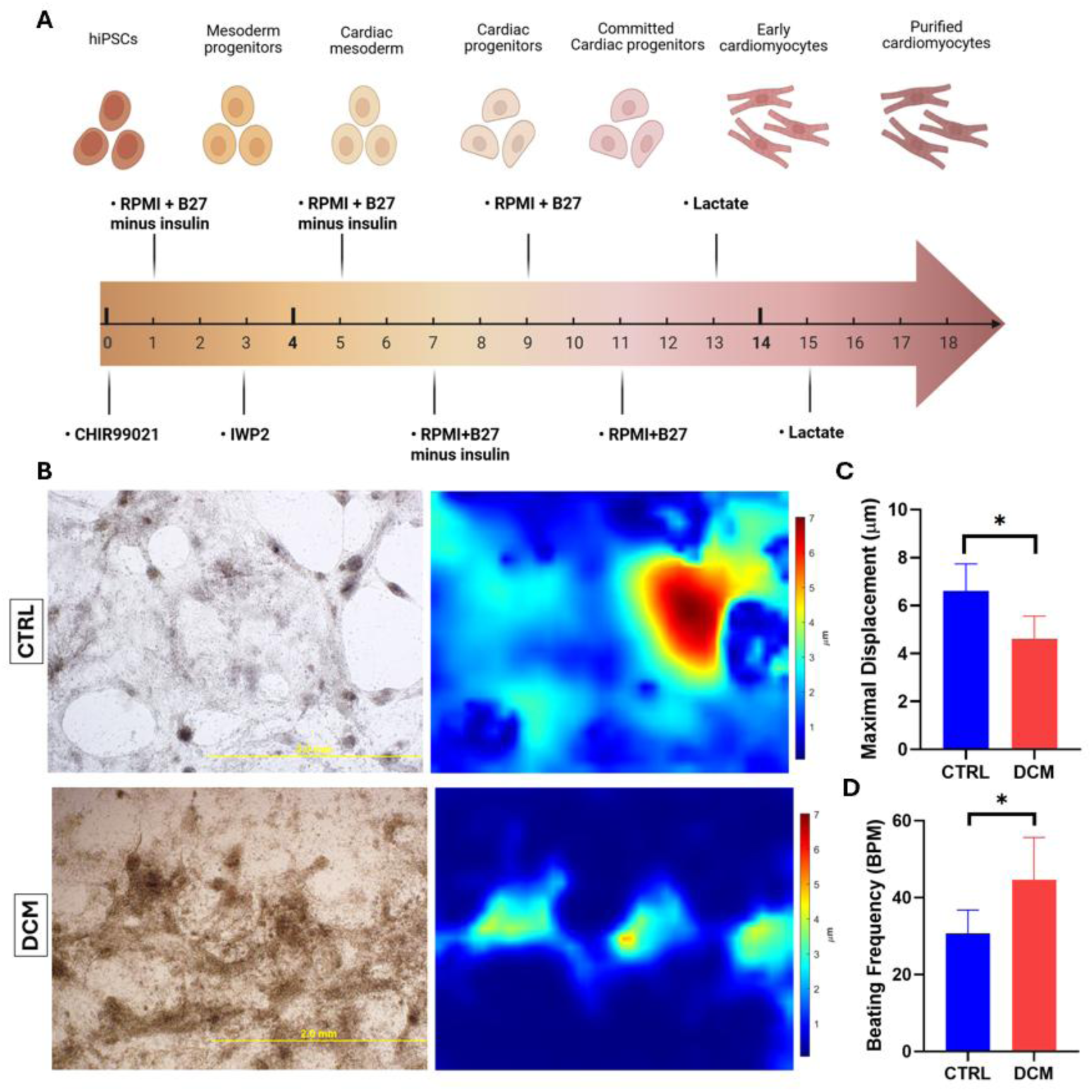
(A) Schematic overview of the differentiation timeline from hiPSCs to CMs, highlighting different developmental stages. (B) Representative images of beating hiPSC-CM monolayers from control and DCM lines, along with corresponding displacement heatmaps. (C) Quantification of maximal displacement and (D) beating frequency of hiPSC-CMs at day 18 post-differentiation (n=6) (* indicates statistical significance at p<0.05).

### 3.3. Differentiation Characterization

The differentiation efficiency was further assessed by IF staining for cardiac-specific biomarkers (**Fig. 4A and S5-7**). Cardiac troponin T (cTnT), which is a CM-specific structural protein and a key component of the sarcomeric contractile apparatus, was expressed in approximately 90% of the cells in both DCM and control lines, demonstrating a high efficiency of CM differentiation (**Fig. 4B**). As cTnT is selectively expressed in CMs, its presence served as a reliable indicator of successful CM differentiation. The differentiated hiPSC-derived CMs (hiPSC-CMs) exhibited a relatively immature, fetal-like cardiac phenotype, characterized by a rounded to slightly polygonal morphology and centrally located nuclei. IF staining for cTnT also confirmed the presence of sarcomeric structures; however, these appeared underdeveloped and less organized, consistent with the structural characteristics of early-stage CMs.

**Fig. 4.**
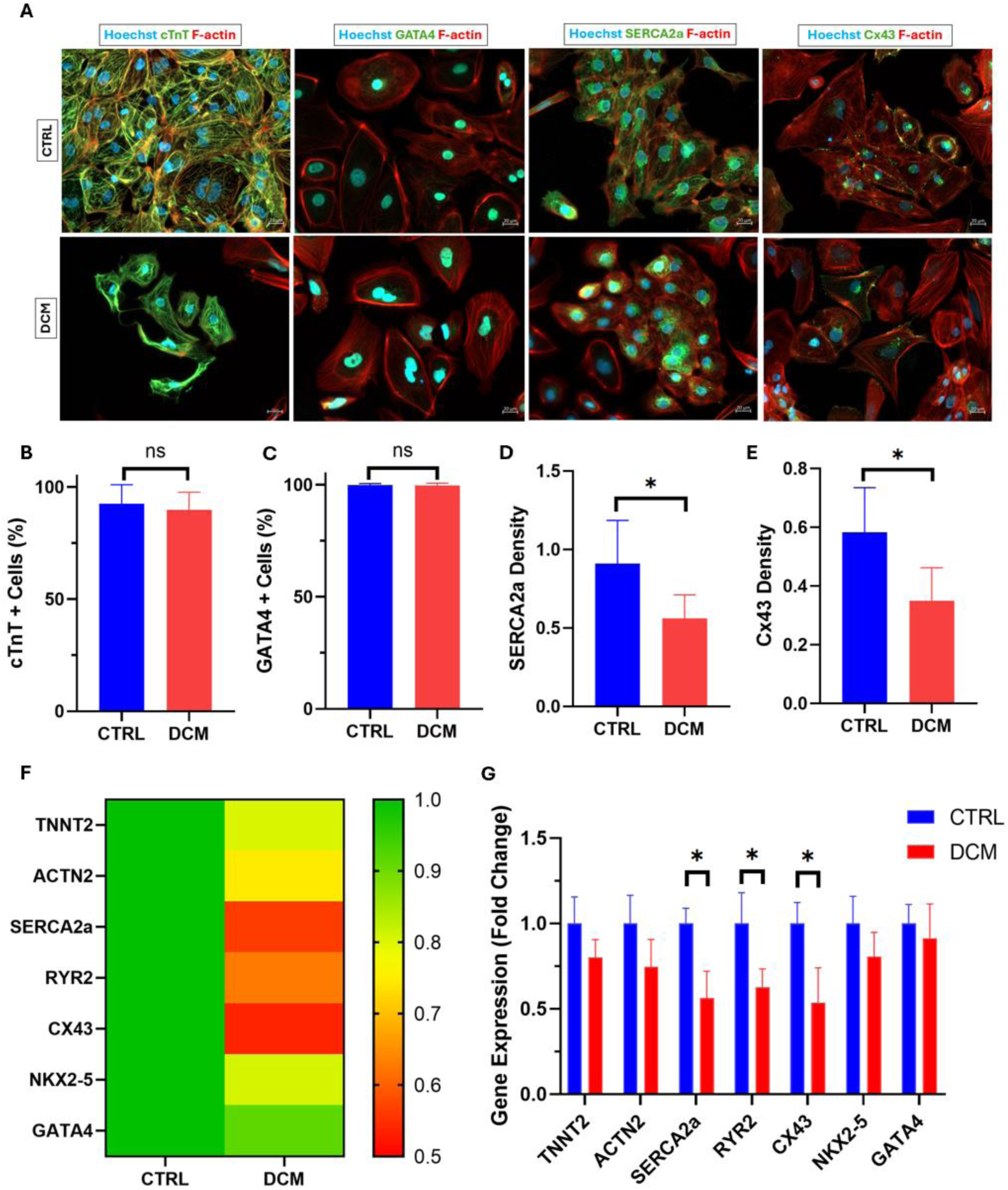
(A) ICC of cardiac-specific markers, showing cTnT, GATA4, SERCA2a, and Cx43, all stained in green. Nuclei and F-actin were shown in blue and red, respectively. Scale bar = 20 μm. (B) Percentage of cTnT-positive cells indicating CM differentiation efficiency. (C) Percentage of GATA4-positive cells demonstrating effective cardiogenic differentiation in both lines. (D) Quantification of SERCA2a and (E) Cx43 densities, showing significant differences between control and DCM lines (n=6). (F) Heatmaps and (G) corresponding graphs of gene expression changes in the DCM group, normalized to the control group (n=6) (ns indicates not statistically significant (p>0.05), and * denotes statistical significance at p<0.05).

In addition, GATA4, which is a transcription factor essential for the initiation of cardiogenic differentiation, was detected in nearly 100% of cells across both groups, confirming robust cardiogenic differentiation (**Fig. 4C**). Furthermore, SERCA2a, an ATP-dependent calcium pump critical for sarcoplasmic reticulum calcium reuptake, and Cx43, a major gap junction protein responsible for intercellular electrical coupling, were positively stained in both lines. These findings indicate the presence of developing calcium-handling machinery and nascent intercellular junctions, both of which are essential for effective excitation-contraction coupling in CMs [25]. Notably, SERCA2a and Cx43 expression levels were quantified from marker density in IF images and were higher in healthy controls (**Fig. 4D-E**). This suggests impaired calcium handling and reduced electrical connectivity in the DCM-derived CMs. Collectively, the characterization results supported successful cardiac differentiation while reflecting the structural immaturity typically associated with hiPSC-CMs at this stage [26].

### 3.4. Gene Expression

Gene expression in control and patient-derived hiPSC-CMs was analyzed by qRT-PCR. The data were calculated as fold changes relative to healthy hiPSC-CMs, normalized to 1, and the resulting heatmaps and graphs are shown in **Fig. 4F-G**. Transcription factors NKX2-5 and GATA4, both of which regulate early cardiac specification and differentiation, showed no statistically significant differences between the control and DCM groups. This finding aligns with the developmental stage of hiPSC-CMs at day 18 post-differentiation, as the principal cardiac transcriptional programs were already activated, and the cells were committed to a CM fate [27]. Similarly, the expression of structural genes TNNT2 (encoding cTnT) and ACTN2 (encoding α-actinin-2) was not significantly altered. This suggests that the core sarcomeric cytoskeleton is relatively preserved in patient-derived CMs, which is expected given that sarcomeric organization typically occurs early during CM differentiation and is less sensitive to disease-specific changes at the transcript level in immature hiPSC-CMs [28].

In contrast, a statistically significant downregulation was observed for Cx43, SERCA2a, and ryanodine receptor 2 (RYR2), all of which play critical roles in calcium handling, electromechanical junctions, and action potential propagation. This reduced expression in the DCM group indicates early impairments in excitation-contraction coupling, consistent with the disease phenotype [29]. Given the relatively immature state of hiPSC-CMs at day 18, defects in calcium cycling and electrical connectivity are expected to be among the earliest detectable functional abnormalities. However, broader structural defects could become evident only at later stages of maturation or at the protein level. Therefore, these results highlight that, while CM differentiation, cardiac lineage specification, and sarcomere organization were mainly preserved in patient-derived hiPSC-CMs, DCM-specific functional deficiencies had already emerged at the molecular level.

### 3.5. Tissue Optimization

To replicate the native microenvironment for encapsulated cardiac cells, we utilized a composite hydrogel comprising fibrin and Geltrex. Fibrin, a natural biopolymer, undergoes rapid polymerization upon the interaction of fibrinogen and thrombin, forming a provisional matrix [30]. Geltrex, a basement membrane extract primarily composed of laminin, collagen IV, entactin, and heparan sulfate, was incorporated to supply essential extracellular matrix (ECM) components, including structural proteins and proteoglycans [31]. Despite its favorable biological properties, fibrin is inherently prone to rapid degradation both *in vivo* (through fibrinolysis) and *in vitro* (due to the activity of plasmin and other proteases secreted by embedded cells) [30, 32]. To mitigate premature matrix degradation and ensure structural integrity during tissue formation, aprotinin, a fibrinolysis inhibitor, was supplemented in the culture medium. This approach has been previously demonstrated to enhance matrix stability and facilitate ECM deposition and remodeling during tissue maturation [33, 34].

To recapitulate DCM *in vitro* and measure contractile forces, we utilized an accessible PDMS-based chip with two muscle chambers, each containing a vertical pillar pair. The device fits into each well of a standard 12-well plate, facilitating high-throughput applications of the HOC platform by enabling the simultaneous formation of 24 microtissues in a single culture plate (**Fig. S8**). The flexible pillars function as structural anchors and support tissue attachment while preserving passive mechanical tension during auxotonic contractions. Their deflection, caused by tissue-generated forces, enables direct quantification of contractility according to the bending beam theory [35]. A linear force-displacement equation was used for the calculation of contractile force, as explained in our previous report [22].

The device was preliminarily validated using primary CMs, demonstrating the formation and compaction of rat EHTs, with spontaneous beating observed from day 3 and sustained through day 7 of culture. The CMs exhibited aligned and elongated morphology, an interconnected troponin network, and dynamic calcium transients. Furthermore, real-time mapping of the integrated beads revealed a progressive increase in contractile robustness over time in these EHTs (**Fig. S9-10**). Neonatal rat CMs have been extensively used as a robust preliminary validation model for human HOC platforms [36–38]. Our results demonstrate that this system holds strong potential for the development of human EHTs.

The chip was further used to fabricate millimeter-scale beating human cardiac tissues composed of differentiated hiPSC-CMs from both DCM and control subjects, embedded in a fibrin/Geltrex ECM. A fraction of HCFs was added to hiPSC-CMs prior to cell encapsulation, owing to their established role in extracellular matrix synthesis and remodeling, which offsets the limited migratory behavior and matrix-remodeling capacity of CMs, whose primary function is to generate contractile force [39–41]. This strategy is anticipated to facilitate tissue assembly and compaction, while enhancing contractile performance by promoting intercellular signaling [42, 43]. Various cell types have been co-cultured with CMs in EHTs to support tissue formation and maturation, including fibroblasts [44], endothelial cells [45], epicardial cells [46], and immune cells [47]. Notably, tissue functionality is strongly influenced by the seeding ratio of CMs to non-CMs [48].

To determine the optimal ratio of CMs to non-CMs for our EHT function, we evaluated three different seeding conditions using hiPSC-CMs and HCFs, including 90:10, 80:20, and 70:30. In each experimental group, the cell mixture was encapsulated in a Fibrin/Geltrex hydrogel with a total density of 4×10^7^ cells/ml and seeded within microfabricated chambers. Over time, the cell–ECM composite compacted into a tissue strip suspended between the pillars, capable of spontaneous contractions that induced pillar deflection.

Tissues seeded with a 90:10 ratio failed to maintain structural integrity over a 14-day culture period, exhibiting incomplete tissue formation, spontaneous detachment from the anchoring pillars, or tissue rupture. This outcome is likely attributable to the insufficient presence of fibroblasts, which are essential for ECM remodeling and structural cohesion during early tissue formation. In contrast, the 80:20 and 70:30 groups both supported robust tissue strip formation within 24 hours. However, spontaneous beating onset and strength varied significantly between these two ratios. The 80:20 group initiated spontaneous contractions by day 4 and maintained consistent, stronger contractions through day 14. In comparison, the 70:30 group exhibited a delayed onset of contractile activity, with beating starting on day 8 and remaining consistently weaker overall. IF staining confirmed the presence of aligned, elongated CMs with organized troponin networks in both groups (**Fig. 5A-B)**, with a higher troponin density observed in the 80:20 group, indicating enhanced cellular maturation and sarcomeric assembly (**Fig. S11-12**).

**Fig. 5.**
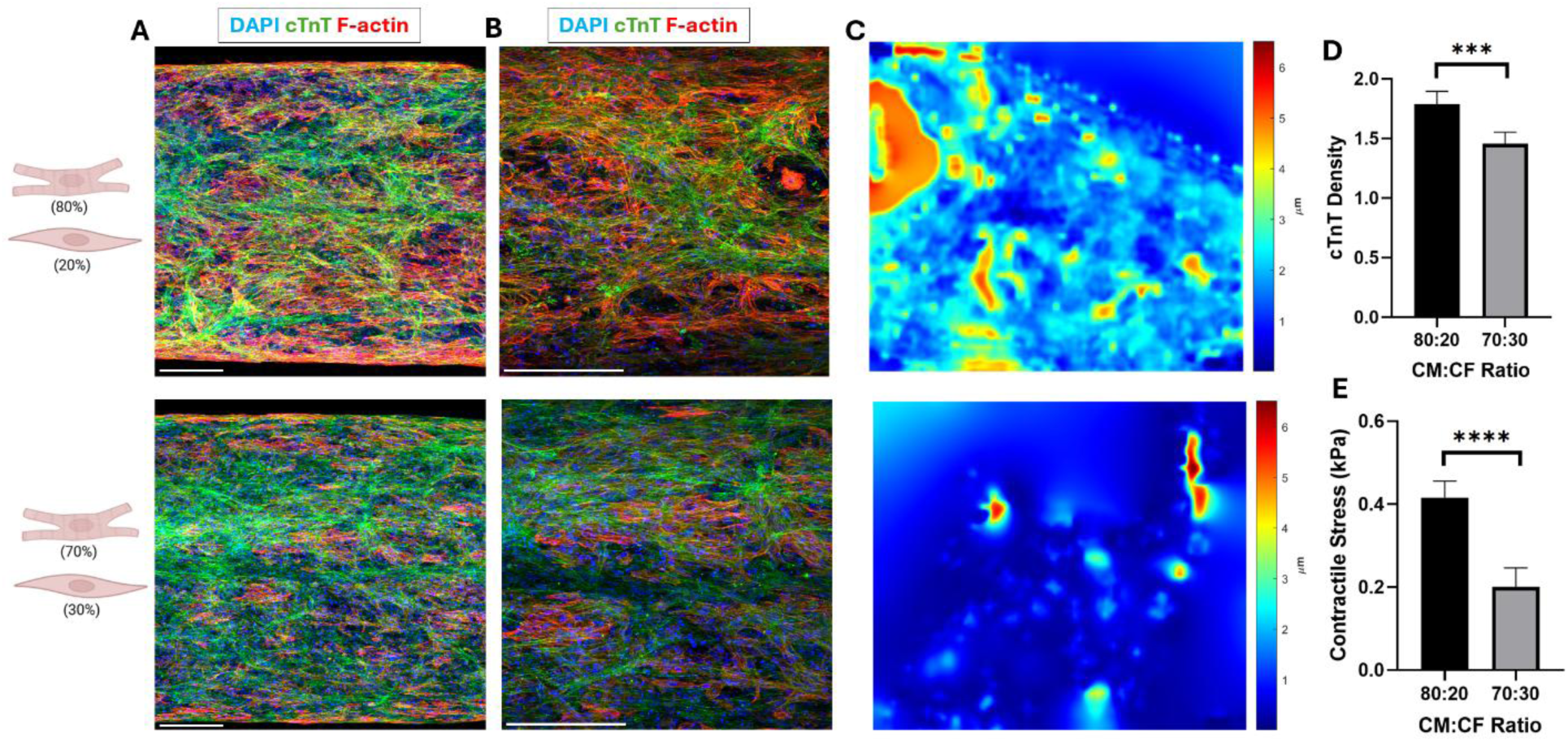
(A) Confocal images of IF-stained EHTs with different CM:CF ratios (80:20 and 70:30), acquired at 10× and (B) 20× magnifications, showing cTnT (green), DAPI (blue), and F-actin (red). Scale bars = 200 μm. (C) Bead displacement maps illustrating local contractile activity across tissues with 80:20 (top image) and 70:30 (bottom image) CM:CF ratios. (D) Quantification of troponin density in EHTs with different cell ratios. (E) Measured contractile stress for EHTs with varying CM:CF ratios (n=6) (*** and **** indicate statistical significance at p<0.001 and p<0.0001, respectively).

To complement the conventional pillar deflection method for contractility assessment, fluorescent microbeads were incorporated into the cell–ECM mixture prior to seeding. These embedded beads enabled real-time mapping of local contractile dynamics across the entire tissue using a custom MATLAB algorithm (**Fig. 5C** and **Movie S3-4**). Quantitative analysis revealed that tissues in the 80:20 group exhibited significantly higher contractile stress throughout the tissue than those in the 70:30 group (**Fig. 5D-E**). Moreover, the 80:20 group demonstrated more uniform bead displacement, indicative of greater contractile homogeneity, resulting in more substantial, more synchronized pillar deflection. In contrast, the 70:30 tissues exhibited regional contractile heterogeneity, with limited bead displacement observed in small areas near the pillar anchors, while the majority of the tissue displayed minimal contractile activity. This spatial imbalance limited whole-tissue force generation and reduced overall pillar deflection, despite successful tissue formation. These findings underscore the importance of an optimized stromal cell fraction for uniform contractile output.

The integration of fluorescent beads into the tissue represents a significant advancement over traditional pillar-only measurements. While pillar deflection provides a global readout of force generation, it is inherently limited to the mechanical response at anchoring points. The bead-based system offers spatially resolved, quantitative insight into local contractile behavior across the tissue, capturing subtle variations in contractility that may be undetectable via pillar displacement alone. Additionally, bead movement proximal to the pillar area can be used to further refine force calculations and validate pillar-based measurements. This real-time bead-tracking system employs a 3D TFM approach, enabling high-resolution contractile mapping that complements and enhances conventional biomechanical assessment methods. Therefore, the 80:20 ratio of hiPSC-CMs to HCFs emerged as the optimal condition, supporting rapid tissue formation, synchronized beating, and enhanced overall contractile performance. This optimized ratio was subsequently used in further experiments for drug screening and disease modeling validations.

### 3.6. Tissue Structure

Control and DCM EHTs were generated by using both control- and patient-derived hiPSCs in combination with HCFs at an optimized 80:20 ratio. Encapsulation within the fibrin/Geltrex hydrogel, followed by seeding into the cardiac chambers, enabled rapid gelation upon thrombin addition. Within 24 hours, cell-mediated matrix remodeling led to the formation of tissue constructs anchored between the flexible PDMS pillars. Over two weeks of culture, progressive tissue compaction was observed, characterized by a reduction in tissue width and increased structural cohesion.

Control EHTs exhibited significantly higher compaction compared to DCM tissues, as evidenced by a smaller tissue width (**Fig. 6A**). This lower compaction suggests impaired ECM remodeling capacity in DCM EHTs, potentially due to intrinsic structural or functional defects associated with the disease phenotype. These findings align with previous reports indicating reduced matrix remodeling ability in DCM tissues, further validating the use of this model for DCM-specific studies [49–51]. Additionally, both control and DCM EHTs formed at consistent heights relative to the PDMS base, confirming reproducibility in tissue positioning and supporting the accuracy of the formulation derived from Microsquisher analysis (**Fig. 6B**). Moreover, robust tissue formation across multiple devices and replicates highlighted minimal batch-to-batch variability (**Fig. 6C**). This high degree of consistency could be attributed to the use of the lactate purification method, which enriched CM purity post-differentiation, and the consistent inclusion of primary HCFs across conditions, both of which are critical factors for reliable ECM remodeling and tissue integrity.

**Fig. 6.**
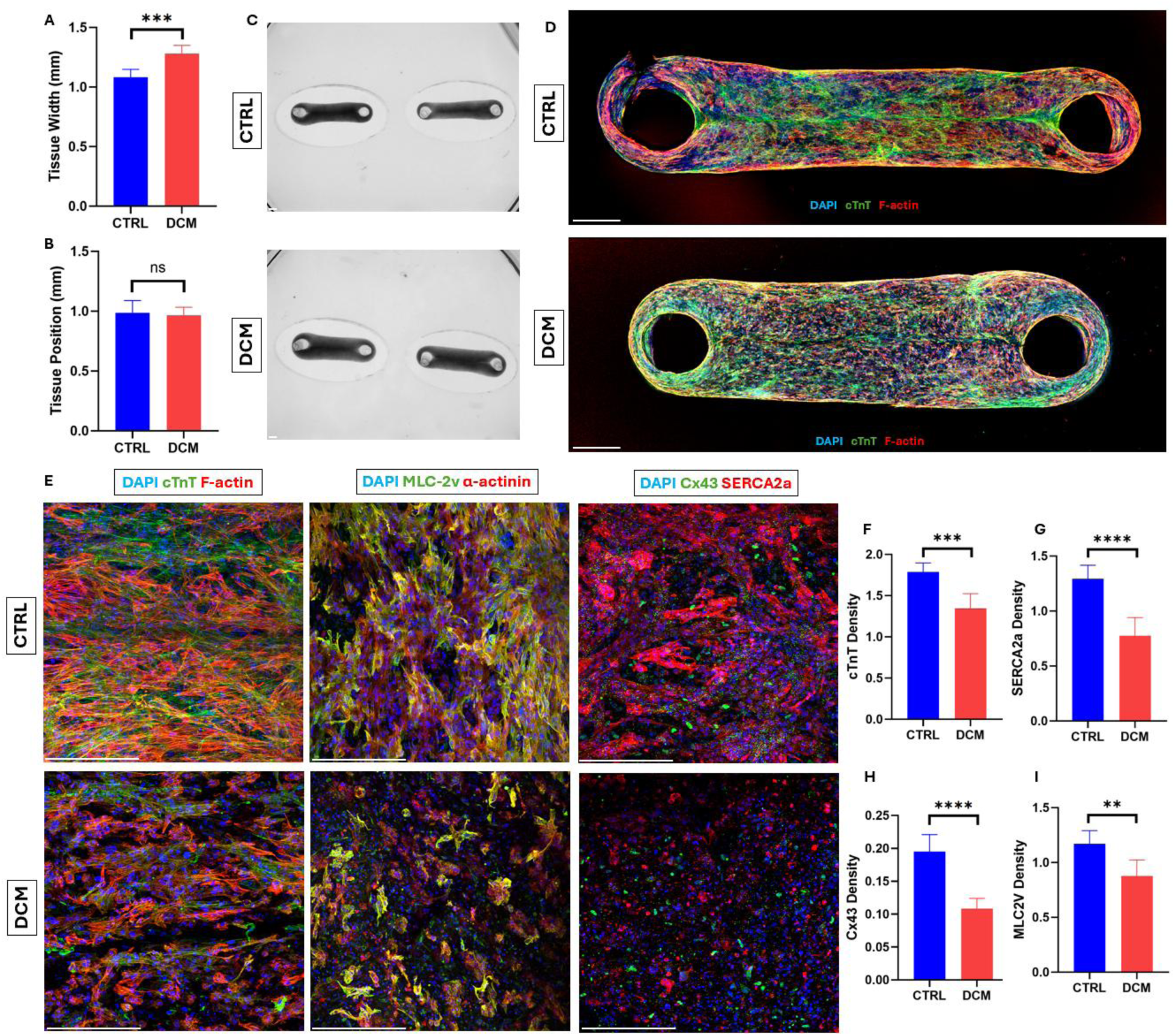
(A) Lateral tissue compaction in control and DCM EHTs after two weeks of culture. (B) Height position of the formed tissues, quantified as the distance from the PDMS base (n=6). (C) Reproducible tissue formation in both control and DCM groups. Scale bars = 500 μm. (D) Confocal images of whole-mount stained control and DCM tissues, showing cTnT (green), DAPI (blue), and F-actin (red). Scale bars = 500 μm. (E) Confocal images of immunofluorescently stained control and DCM EHTs, acquired using a 20x objective. Left: cTnT (green), DAPI (blue), and F-actin (red); Middle: MLC-2v (green), DAPI (blue), and α-actinin (red); Right: Cx43 (green), DAPI (blue), and SERCA2a (red). Scale bars = 200 μm. (F–I) Quantification of marker density for (F) cTnT, (G) SERCA2a, (H) Cx43, and (I) MLC-2v in control and DCM groups (n=6) (ns indicates not statistically significant (p>0.05), and **, ***, and **** denote statistical significance at p<0.01, p<0.001, and p<0.0001, respectively).

To further evaluate tissue structure and cell distribution within the EHTs, whole-mount immunostaining was performed after two weeks of culture. Samples were co-stained with DAPI, phalloidin, and cTnT to visualize nuclear localization, cytoskeletal organization, and CM-specific structure, respectively (**Fig. S13**). Representative confocal images of CTRL and DCM tissues are shown in Fig. 6D, illustrating the overall morphology and structural integrity of the constructs, with a uniform distribution of CMs within EHTs. This homogeneous cell dispersion and consistent organization reflect the successful encapsulation of high-density CMs within the ECM. Also, DCM tissues exhibited reduced compaction, while control tissues displayed a more aligned, elongated, and interconnected CM network, underscoring the structural deficits associated with the disease [52].

Furthermore, high-resolution IF imaging was performed to evaluate structural and cellular organization using cTnT/Phalloidin, SERCA2a/Cx43, and MLC-2v/α-actinin staining, as shown in **Fig. 6E**. In control tissues, CMs appeared well-aligned and elongated, displaying an organized troponin complex with a clear striated pattern and well-defined contractile units (**Fig. S14**). In contrast, DCM tissues exhibited disorganized cellular alignment, reduced elongation, and disrupted troponin organization, consistent with the structural and functional impairments associated with the DCM phenotype [53]. Additionally, control tissues demonstrated elongated and striated sarcomeres and strong MLC-2v expression, indicative of mature ventricular identity and sarcomere integrity. Conversely, DCM tissues showed disrupted MLC-2v patterns, reduced sarcomere alignment, and diminished elongation, reflecting an immature and disorganized contractile architecture (**Fig. S15**). SERCA2a expression was also notably stronger and more localized in control tissues, suggesting efficient calcium handling. In DCM tissues, however, SERCA2a appeared diffuse and attenuated, suggesting impaired calcium reuptake. In parallel, Cx43 was distinctly localized at intercellular junctions in control EHTs, forming punctate structures that signify functional gap junctions and coordinated electrical coupling. On the other hand, Cx43 was fragmented and dispersed in DCM EHTs, suggesting compromised intercellular communication (**Fig. S16**).

Quantification of IF images confirmed significantly higher densities of cTnT, SERCA2a, Cx43, and MLC-2v in control tissues, reflecting enhanced sarcomeric organization, effective calcium cycling, robust electrical connectivity, and a mature ventricular phenotype (**Fig. 6F-I**). Furthermore, the ratio of MLC-2v to α-actinin intensity was evaluated as a marker of ventricular differentiation efficiency, given that MLC-2v is specific to ventricular CMs, whereas α-actinin is expressed across all CM subtypes [54]. This analysis revealed a high ventricular differentiation efficiency (∼90%) in both groups, consistent with the expected outcomes of the applied ventricular differentiation protocol (**Fig. S17**).

### 3.7. Drug Screening

To evaluate the drug screening capability of our HOC model, we investigated the functional response of both control and DCM tissues to norepinephrine (NE), which is a well-established β-adrenergic agonist known to enhance the cardiac contractile force and beating rate [55, 56]. For NE stimulation, a concentration of 100 µM was selected based on previous studies on EHTs [14, 22, 23]. By monitoring contractile activity and calcium dynamics before and after NE treatment, we aimed to determine whether the model could accurately reproduce pharmacological effects in a physiologically relevant manner (**Movie S5-8**).

Following NE treatment, control tissues exhibited a robust and expected response. As shown in **Fig. 7A-C**, NE stimulation significantly elevated peak contractile force, increased contractile stress, and accelerated the spontaneous beating frequency of the tissues. Concurrently, calcium transient amplitude also increased, indicative of improved intracellular calcium cycling **(Fig. 7D-F**). These outcomes are consistent with the known mechanisms of β-adrenergic stimulation, which include elevated calcium influx via upregulation of L-type calcium channels, enhanced sarcoplasmic reticulum (SR) calcium uptake via phospholamban phosphorylation, and increased excitation-contraction coupling efficiency [57, 58]. These results collectively confirm the functional responsiveness of the constructed human EHTs, validating the platform’s physiological relevance.

**Fig. 7.**
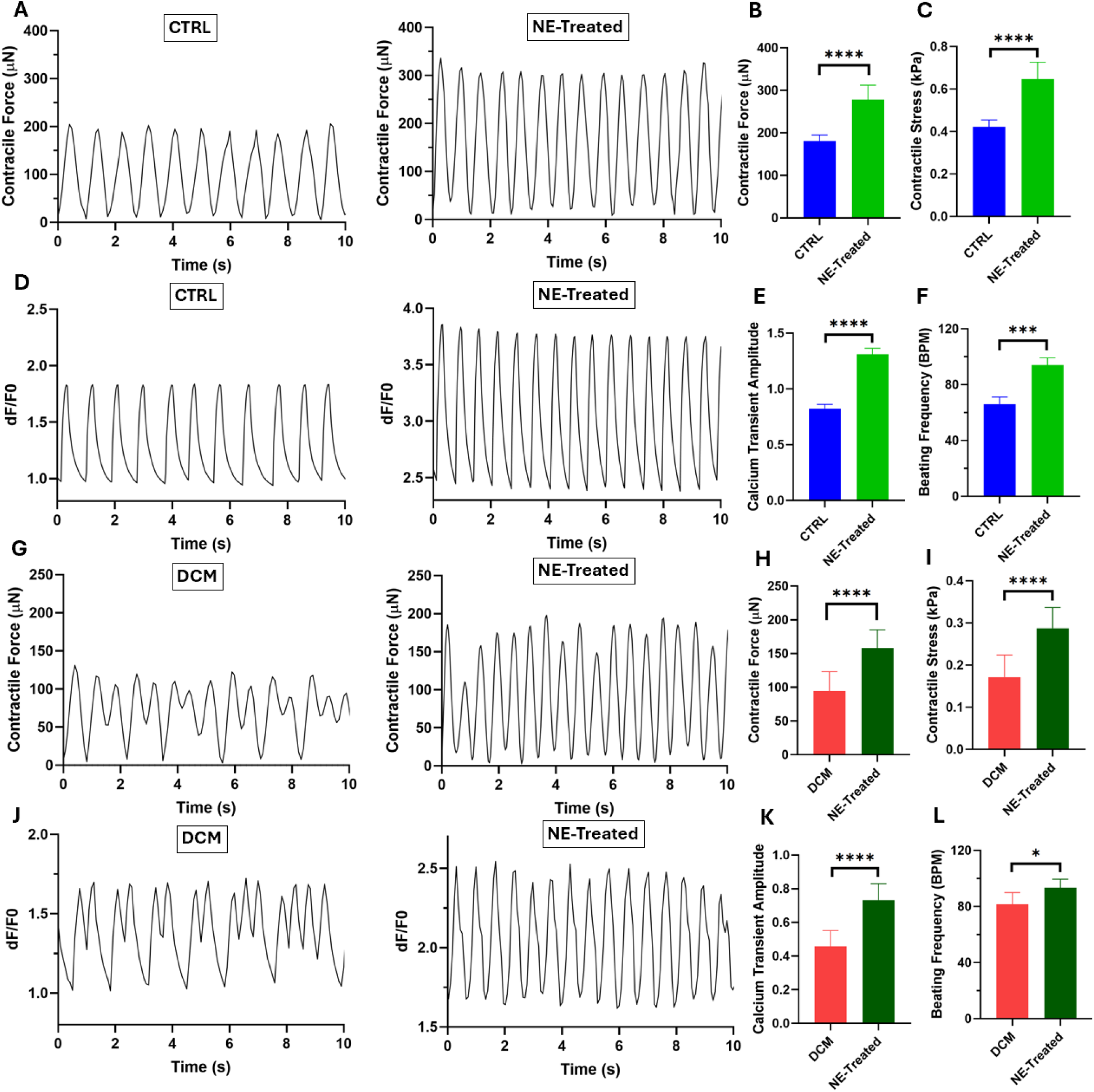
(A) Beating patterns of control (CTRL) and norepinephrine (NE)-treated control EHTs, with quantification of (B) contractile force and (C) contractile stress. (D) Calcium transient traces of CTRL and NE-treated control EHTs, with quantification of (E) calcium transient amplitude and (F) beating frequency. (n=6) (G) Beating patterns of DCM and NE-treated DCM EHTs, with quantification of (H) contractile force and (I) contractile stress. (J) Calcium transient traces of DCM and NE-treated DCM EHTs, with quantification of (K) calcium transient amplitude and (L) beating frequency. (n=6) (*, *** and **** indicate statistical significance at p<0.05, p<0.001 and p<0.0001, respectively).

Moreover, DCM tissues displayed functional hallmarks of the disease, including reduced baseline contractility and irregular, arrhythmic beating patterns, indicative of impaired calcium handling and immature excitation-contraction coupling. This marked electrophysiological instability was indicated by irregular spontaneous beating and the presence of early afterdepolarizations (EADs), inducing episodes of ventricular tachycardia (**Fig. 7G,J**). These arrhythmic patterns are often triggered by delayed repolarization and intracellular calcium overload, resulting in abnormal depolarizing currents during the repolarization phase [59]. In DCM, such disturbances are commonly attributed to impaired SR calcium reuptake, leaky ryanodine receptors, or dysregulated sodium-calcium exchanger (NCX) activity, promoting reentrant arrhythmias [60, 61].

Following NE stimulation, significant increases in calcium transient amplitude, contractile force, contractile stress, and beating frequency were also observed in DCM tissues, indicating β-adrenergic responsiveness (**Fig. 7H,I,K**). Interestingly, DCM tissues exhibited a marked reduction in EAD-induced arrhythmic events despite an overall increase in beating rate, indicating intensification of tachycardia (**Fig. 7L**). This partial improvement in beating patterns may be attributed to enhanced calcium handling and repolarization dynamics mediated by β-adrenergic signaling [62]. Particularly, increased SR calcium uptake, along with more synchronized calcium release from intracellular stores, likely contributed to the observed regularization of contractile activity [63]. These findings suggest that in our partially mature DCM model, β-adrenergic stimulation may exert a compensatory effect, transiently restoring electrical and mechanical coordination despite underlying deficits in calcium homeostasis.

### 3.8. Disease Modeling

In the next step, the functionality of EHTs formed from control and DCM-derived stem cells was compared, revealing distinct differences in contraction and calcium-handling dynamics. DCM tissues exhibited a significantly higher spontaneous beating frequency than controls, consistent with the occurrence of ventricular tachycardia (**Fig. 8A**). This arrhythmic behavior in DCM EHTs can be attributed to impaired electrical coupling and disrupted calcium homeostasis, hallmarks of dilated cardiomyopathy. The observed EAD-induced ventricular tachycardia in DCM EHTs can be mechanistically explained by the significant downregulation of key calcium-handling and electrical coupling genes, as revealed by our qRT-PCR analysis. Specifically, DCM tissues showed reduced expression of SERCA2a, CX43, and RYR2. The diminished SERCA2a expression likely contributed to impaired calcium clearance and prolonged cytosolic calcium transients, promoting arrhythmogenic afterdepolarizations. Concurrently, reduced CX43 levels would impair cell-cell gap junctional coupling and electrical propagation, leading to conduction delays and reentry circuits, while decreased RYR2 expression may have exacerbated abnormal calcium release events, further promoting arrhythmic behavior. Together, these molecular defects establish a substrate conducive to EADs and ventricular tachycardia in DCM tissues, consistent with our functional results.

**Fig. 8.**
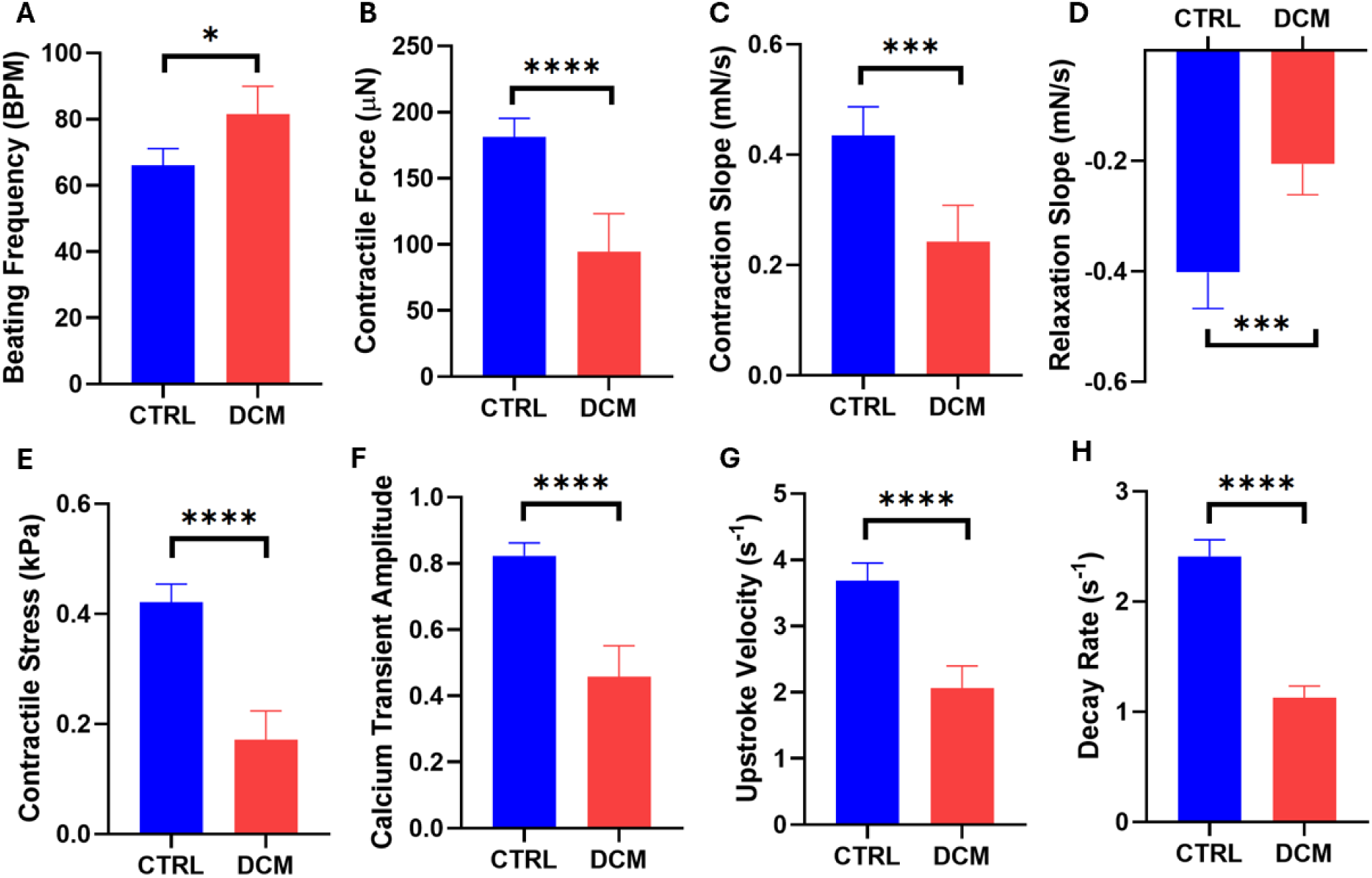
Comparison of functional parameters between DCM and control EHTs, including (A) beating frequency, (B) contractile force, (C) contraction slope, (D) relaxation slope, (E) contractile stress, (F) calcium transient amplitude, (G) upstroke velocity, and (H) calcium decay rate. (n=6) (*, *** and **** indicate statistical significance at p<0.05, p<0.001 and p<0.0001, respectively).

Moreover, control tissues demonstrated significantly superior contractile performance, including higher contractile force (**Fig. 8B**), contraction slope (**Fig. 8C**), and relaxation slope (**Fig. 8D**), indicative of stronger and more synchronized contraction and relaxation cycles. Importantly, the contractile force was normalized to the cross-sectional area of the tissues to calculate the contractile stress, which is a critical parameter for standardization and meaningful comparison across studies (**Fig. 8E**). Also, reporting contractile stress rather than raw force allows for size-independent evaluation of tissue functionality [64]. DCM tissues exhibited significantly lower contractile stress than control tissues, with differences more pronounced than those observed in contractile force measurements, due to reduced tissue compaction in the diseased constructs. Our measured contractile stresses were within the range of previous reports on human EHTs [65–67], however, these values remained lower than in some studies that applied advanced maturation strategies, such as electrical stimulation, mechanical loading, and dynamic and prolonged culture, which promote more mature cardiac phenotypes [39, 68, 69]. On the other hand, our platform, while recapitulating key features of patient-specific disease, still represents a partially matured model, as the measured contractile stress remained markedly lower than that of adult human myocardium (∼44 kPa) [68]. Importantly, the significant functional deficiencies observed between DCM and control tissues highlight the capability of our system in modeling disease-specific phenotypes.

Furthermore, calcium handling properties were markedly enhanced in control tissues, as evidenced by higher calcium transient amplitude (**Fig. 8F**), faster calcium upstroke velocity (**Fig. 8G**), and accelerated calcium decay rate (**Fig. 8H**). These results suggest that control EHTs possess more efficient excitation-contraction coupling mechanisms, enabling more robust and rapid contraction and relaxation dynamics. These defects in contractility and calcium handling have also been reported in recent studies on 3D *in vitro* cardiomyopathy models [51, 70]. The physiological basis for these impairments could be attributed to disrupted structural organization, downregulation of key calcium-handling proteins, and impaired electrical connectivity in DCM tissues, which collectively contribute to deficient contractile performance.

The functional trend in our 3D model aligns with our 2D results for patient-specific iPSC-CMs post-differentiation, in which DCM-derived cells showed increased beating frequency and diminished contractile strength compared to controls. However, the differences between healthy and diseased phenotypes were even more pronounced in the 3D EHT model. In 2D, cells often exhibit immature and less organized behavior, partly due to the lack of appropriate mechanical and structural cues, which can obscure subtle pathological differences [71]. However, 3D constructs provide a biomimetic environment that better recapitulates the native cardiac microarchitecture, cell-cell and cell-matrix interactions, and mechanical load experienced by CMs *in vivo* [72]. Importantly, the 3D HOC model enabled quantification of physiological responses, such as contractile force and stress, that cannot be measured in 2D monolayers [73]. Moreover, the 3D system preserved critical functional hallmarks of patient-specific CMs, including arrhythmic beating patterns in DCM tissues, thereby offering a more faithful disease model. Additionally, the spontaneous beating frequency observed in the 3D constructs was higher than in 2D cultures and fell within a range of the human heart in vivo (∼ 60-100 bpm) [74, 75]. This indicates that the 3D environment supports more physiologically relevant pacing activity, likely due to improved cell alignment, mechanical loading, and maturation stimuli provided by the tissue construct.

Furthermore, our study benefits from the standardized, reproducible conditions provided by a commercially available device. Commercial production ensures consistent manufacturing and uniform properties, minimizing batch-to-batch variability and enhancing reproducibility across experiments. This is particularly difficult to achieve with custom microfabricated chips, which often introduce subtle technical variations. The ready-to-use format also lowers the barrier to adoption for other laboratories, enabling straightforward implementation without microfabrication expertise, and supporting broader dissemination and cross-laboratory comparability.

Despite the significant advantages of our model, several limitations remain to be addressed. First, the current platform relies on hiPSC-cardiomyocytes derived from a single DCM patient line, which does not capture the inter-patient variability or the broad spectrum of DCM etiologies, including familial versus idiopathic forms and mutation-specific differences. Although the patient and control subjects were matched on background information, the present findings remain a proof-of-concept single-patient case study. Future studies should incorporate multiple patient-specific lines representing diverse genetic backgrounds to strengthen phenotype discrimination and enhance its relevance for precision medicine. In addition, while our EHTs demonstrate β-adrenergic responsiveness, pharmacological evaluation was restricted to norepinephrine at a single concentration (100 μM). As such, our results support an initial validation of pharmacological sensitivity rather than a comprehensive drug-screening or safety-efficacy assessment; broader dose-dependent and multi-drug testing will be required to position the platform within preclinical screening pipelines. Furthermore, the absence of patient-derived fibroblasts and other non-myocyte cell types limits our ability to model patient-specific stromal remodeling and its contribution to DCM progression. Future iterations should incorporate additional cardiac cell types to better recapitulate the native myocardium’s multicellular complexity. Finally, the addition of electrical pacing or programmable stimulation modules may further refine the resolution of contractile phenotypes and improve the detection of genotype-dependent functional differences. Moving forward, combining multi-patient cohorts, expanding pharmacological panels, and integrating genomic and clinical data will advance this system toward a robust platform for personalized disease modeling and pharmacological response assessment.

## 4. Conclusion

In summary, our 3D HOC model successfully recapitulated key pathological hallmarks of DCM and demonstrated an anticipated pharmacological response, demonstrating its robust potential for both disease modeling and drug screening applications. The platform reliably reproduced functional differences between control and DCM tissues, including disrupted structural organization of the CM network, reduced contractile stress, and impaired calcium handling in DCM constructs. These results were mechanistically supported by the downregulation of essential calcium-handling and electrical coupling genes. Notably, the response of both groups to β-adrenergic stimulation with NE validated the physiological relevance of the model, with both groups exhibiting the expected increase in contractile force and frequency. These findings collectively underscore the advantages of 3D EHTs over traditional 2D monolayers, particularly in capturing subtle disease phenotypes and enabling quantification of contractile function. By maintaining patient-specific features and enabling real-time functional analysis, this HOC model offers a physiologically relevant and scalable platform for personalized medicine, drug safety and efficacy assessments, mechanistic studies for cardiac pathophysiology, and future integration into high-throughput preclinical screening pipelines.

## Declaration of Competing Interest

M.A. is a co-founder of eNUVIO Inc. All other authors declare no competing financial interest.

## Acknowledgment

Our work is supported by the Natural Sciences and Engineering Research Council of Canada (NSERC) (NSERC, RGPIN-2021-03960, DGECR-2021-00337), Fonds de Recherche du Québec Santé (FRQS) (Chercheurs-boursiers J1 (313837)), Establishment of Young Investigators (324277), Montréal TransMedTech Institute (iTMT), CRCHU Sainte-Justine (CRCHUSJ), and the University of Montréal. We thank Jacques Courtois for his generous donation in support of the Heart-in-a-Dish research project. A.M. gratefully acknowledges the FRQS Doctoral Scholarship, the Merit Scholarship from the Faculty of Medicine of the University of Montréal, and the Michel Bergeron Scholarship from the Department of Pharmacology and Physiology of the University of Montréal. The authors also thank N. Kachurina (RI-MUHC) for technical support during experiments.

## Notes

### Competing Interest Statement

Mark Aurousseau is a co-founder of eNUVIO Inc. All other authors declare no competing financial interest.

