## Supporting Information for "Patient-Specific 3D Heart-On-a-Chip Model of Dilated Cardiomyopathy with Embedded Bead-Based Mapping of Tissue Contractility"

### Supplementary Materials and Methods

**Neonatal Rat Cardiomyocyte Isolation:** Ventricular CMs were extracted from Sprague-Dawley rat neonates (Charles River Laboratories, Canada) with a Neonatal Heart Dissociation Kit (Miltenyi Biotec, Germany). The procedure was approved by the Animal Ethics Committee of the Centre Hospitalier Universitaire Sainte-Justine Research Centre (CRCHUSJ, protocol number: 2023-4833) and complied with the guidelines set by the Canadian Council on Animal Care (CCAC) and the US National Institutes of Health Guide for the Care and Use of Laboratory Animals. Briefly, 3-day-old neonatal rats were euthanized via decapitation, and their hearts were instantly harvested in cold PBS. The ventricles were excised, minced into small fragments, and washed twice with cold PBS. Tissue dissociation was performed using a combination of mechanical and enzymatic methods with a gentleMACS Dissociator (Miltenyi Biotec, Germany), according to the manufacturer's instructions. After dissociation, red blood cells (RBCs) were removed using an RBC lysis solution (Miltenyi Biotec, Germany) based on the manufacturer's recommended protocol. The resulting cell suspension was pre-plated in DMEM/F12 culture medium supplemented with 10% FBS and 1% Pen Strep. After 1 h, the suspended CMs were separated from adherent CFs, which remained attached to the flask. The CMs were then collected via centrifugation at 600 × g for 5 minutes and were immediately prepared for encapsulation by resuspending the cell pellets in the hydrogel solution.

**Rat EHT Formation:** A similar procedure (as described in Section 2.7) was used for rat EHT formation using neonatal rat CMs at a density of 10 million cells/ml. The rat EHTs were cultured in Claycomb medium supplemented with 10% FBS, 1% Pen Strep, 1% GlutaMAX, and 10 µg/mL aprotinin.

### Supplementary Tables

**Table S1.** List of CM primers used in this study

| Gene Symbol | Full Name | Forward Sequence | Reverse Sequence |
|---|---|---|---|
| TNNT2 | Cardiac Troponin T | GGAGGAGTCCAAACCAAAGCC | TCAAAGTCCACTCTCTCTCCATC |
| ACTN2 | α-actinin | CAAACCTGACCGGGGAAAAAT | CTGAATAGCAAAGCGAAGGATGA |
| SERCA2a (ATP2A2) | Sarcoplasmic/endoplasmic reticulum $Ca^{2+}$ ATPase 2a | GGACTTTGAAGGCGTGGATTGTG | CTCAGCAAGGACTGGTTTTCGG |
| RYR2 | Ryanodine Receptor 2 | CATCGAACACTCCTCTACGGA | GGACACGCTAACTAAGATGAGGT |
| CX43 (GJA1) | Connexin 43 | CAATCTCTCATGTGCGCTTCT | GGCAACCTTGAGTTCTTCCTCT |
| NKX2-5 | NK2 Homeobox 5 | AAGTGTGCGTCTGCCTTTCC | TCTTTTCGGCTCTAGGGTCCTT |
| GATA4 | GATA Binding Protein 4 | AATGCCTGCGGCCTCTACA | AGATTTATTCAGGTTCTTGGGCTTC |
| GAPDH | Glyceraldehyde-3-Phosphate Dehydrogenase | GTCTCCTCTGACTTCAACAGCG | ACCACCCTGTTGCTGTAGCCAA |

**Supplementary Figures**

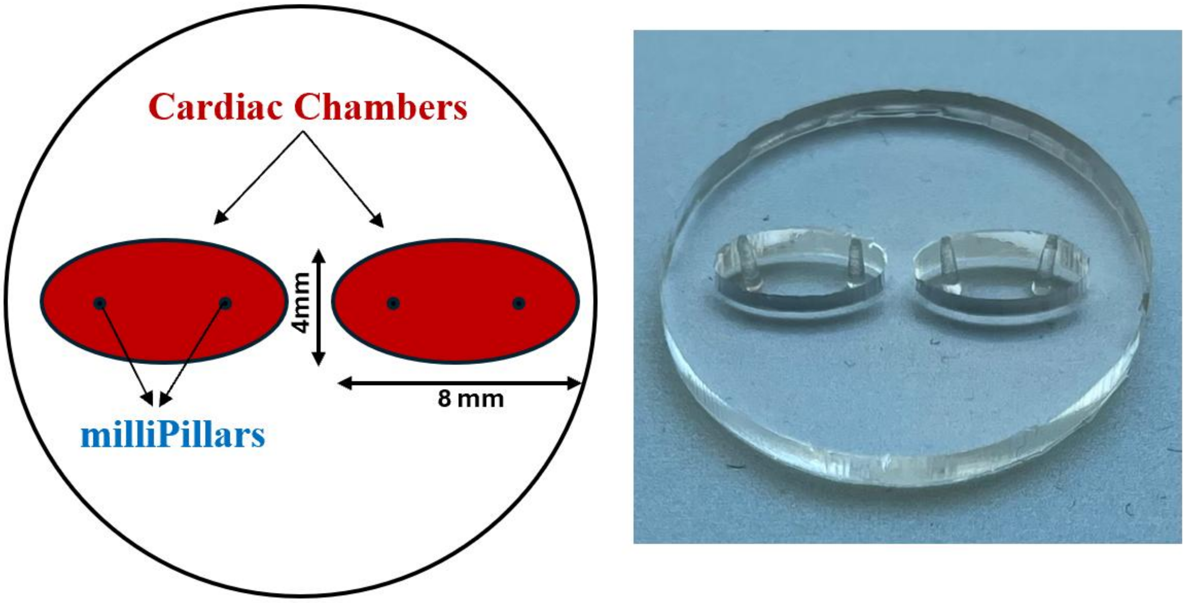


**Fig. S1.** Schematic and actual top-view images of the chip showing its design and chamber dimensions.

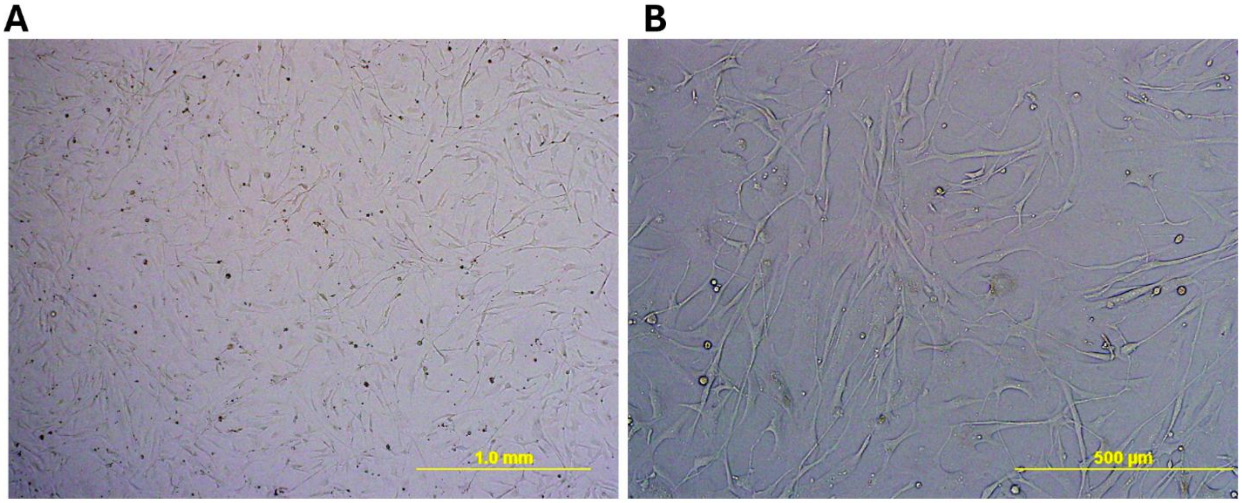


**Fig. S2.** Representative images of HCFs in culture captured under an optical microscope at (A) 4x and (B) 10x magnifications.

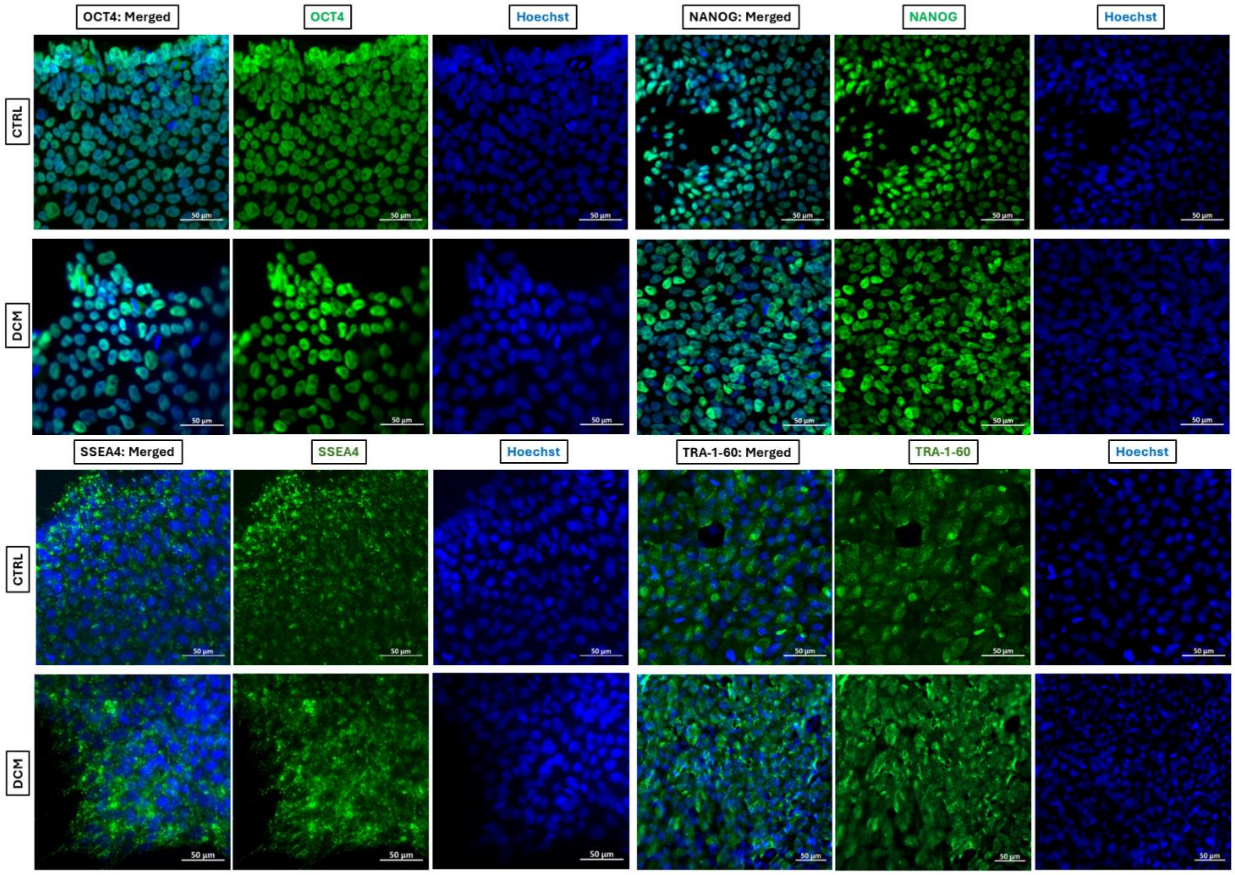


**Fig. S3**. Confocal images showing different channels and merged images of IF staining for main pluripotency markers, including OCT4, NANOG, SSEA4, and TRA-1-60, all depicted in green. Blue channel represents Hoechst. Scale bar = 50 μm.

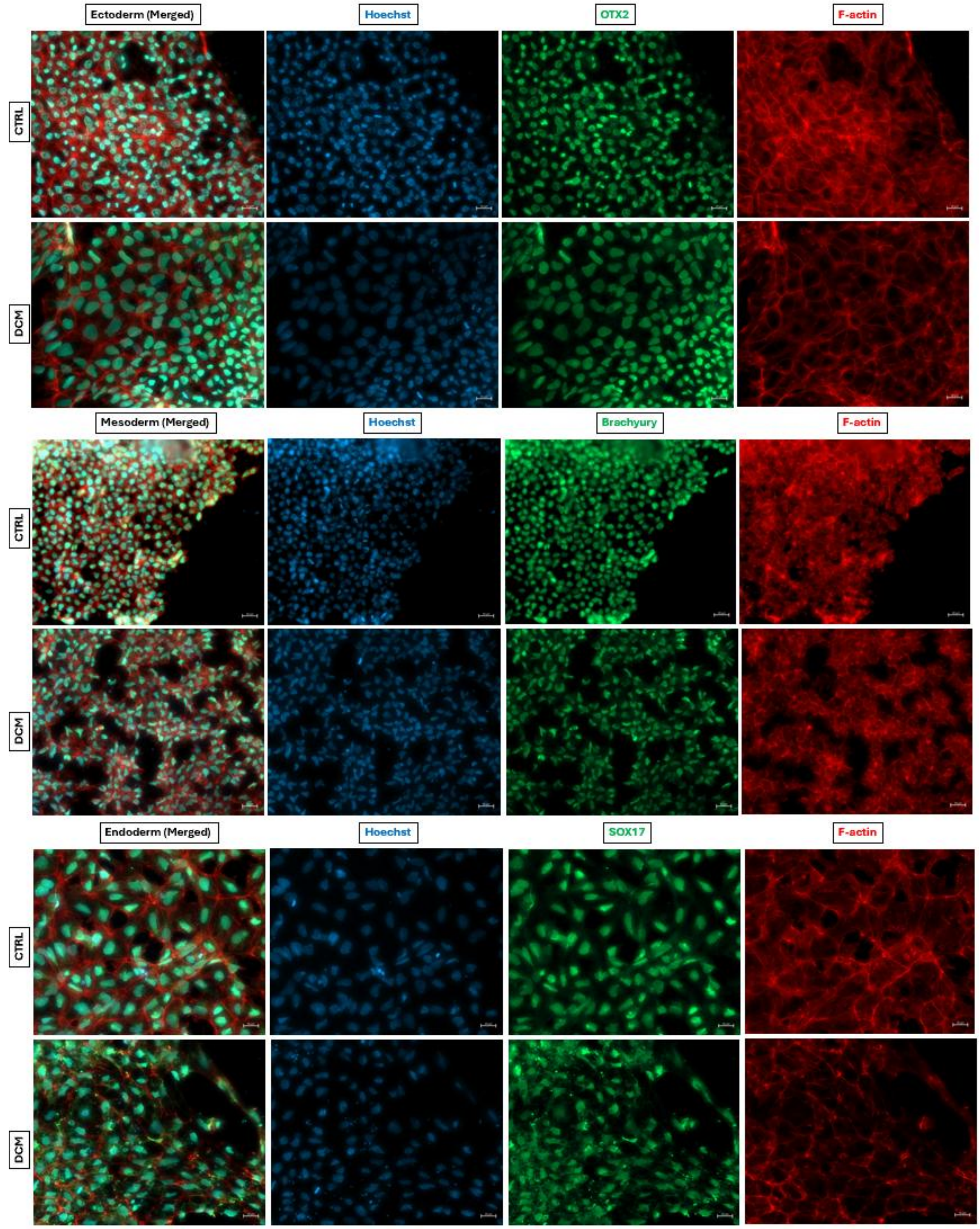


**Fig. S4.** Confocal images showing individual channels and merged images of hiPSC-derived tri-lineage differentiation into ectoderm (OTX2), mesoderm (Brachyury), and endoderm (SOX17), shown in green. Nuclei and F-actin were shown in blue and red, respectively. Scale bar = 20 μm.

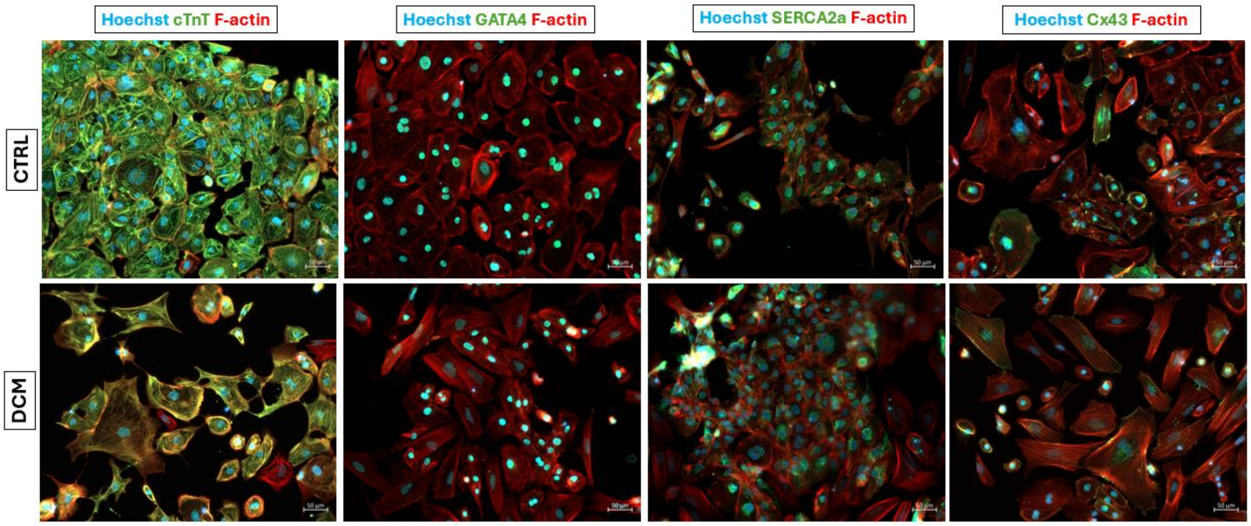


**Fig. S5.** ICC of cardiac-specific markers, demonstrating cTnT, GATA4, SERCA2a, and Cx43, all visualized in green. Nuclei and F-actin were exhibited in blue and red, respectively. Scale bar = 50 µm.

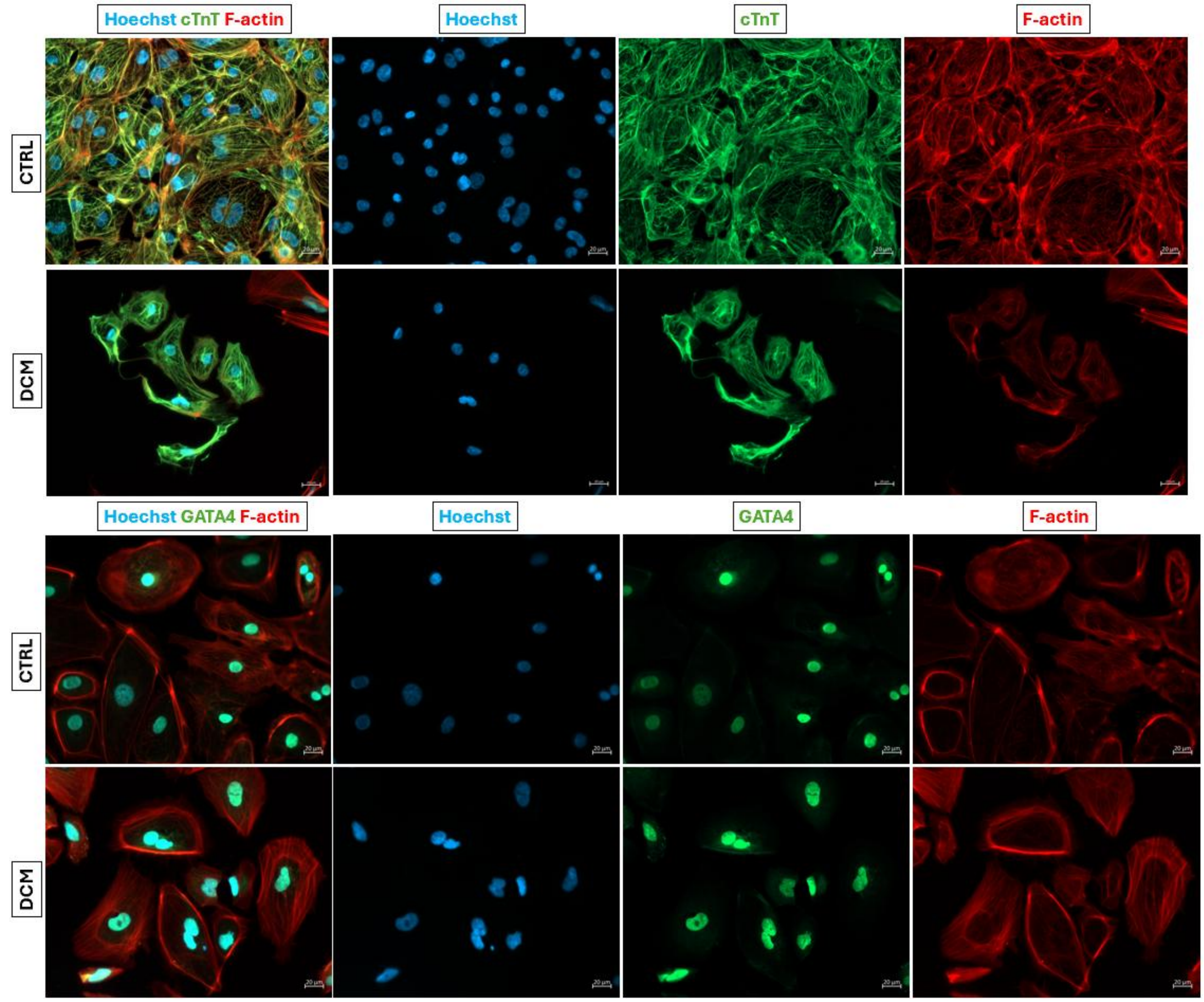


**Fig. S6.** Confocal images illustrating different channels and merged images of IF staining for cardiac-specific markers, including cTnT and GATA4, both indicated in green. Nuclei and F-actin were stained in blue and red, respectively. Scale bar = 20 μm.

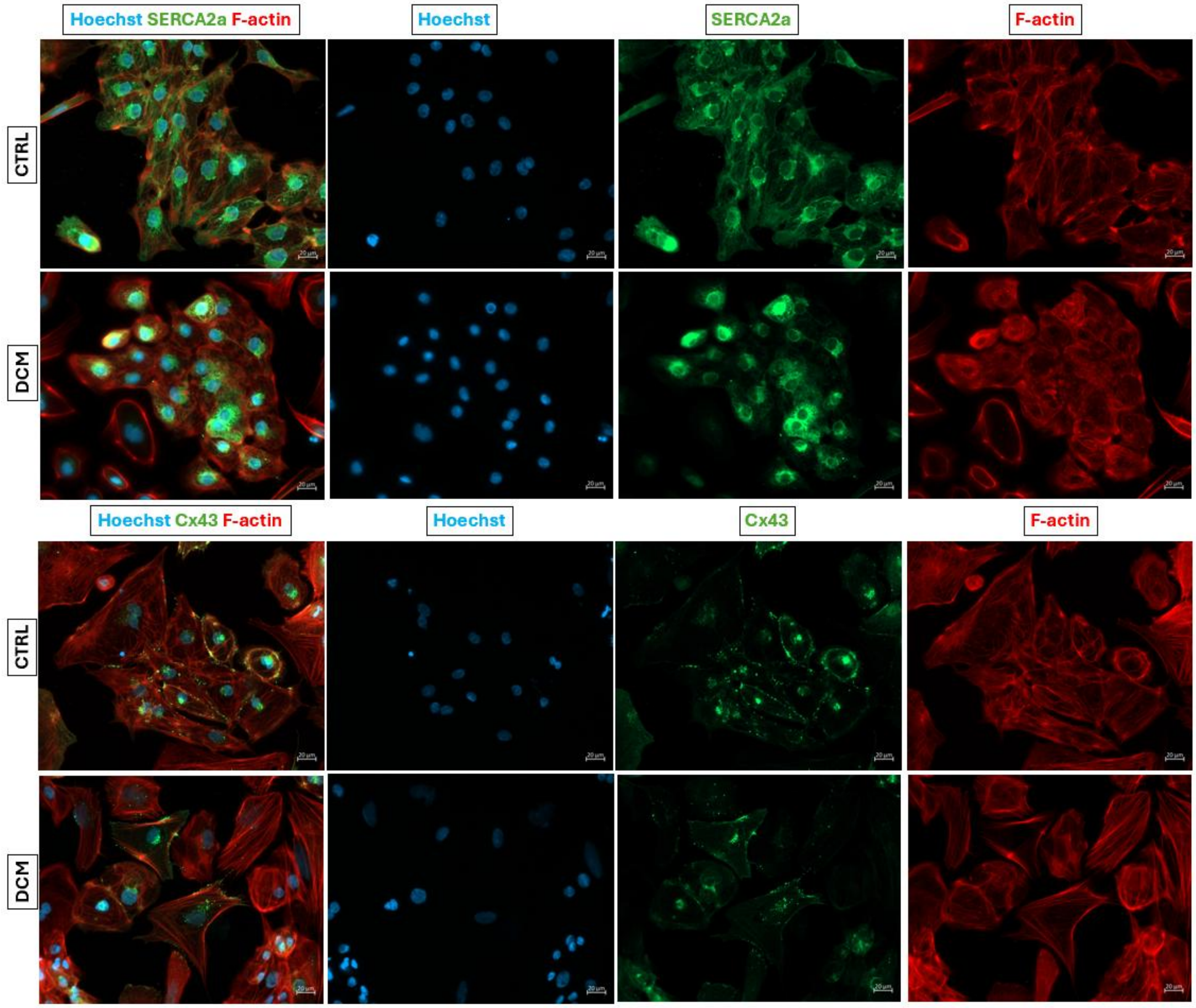


**Fig. S7.** Confocal images illustrating different channels and merged images of IF staining for cardiac-specific markers, including SERCA2a and Cx43, both exhibited in green. Nuclei and F-actin were shown in blue and red, respectively. Scale bar = 20 μm.

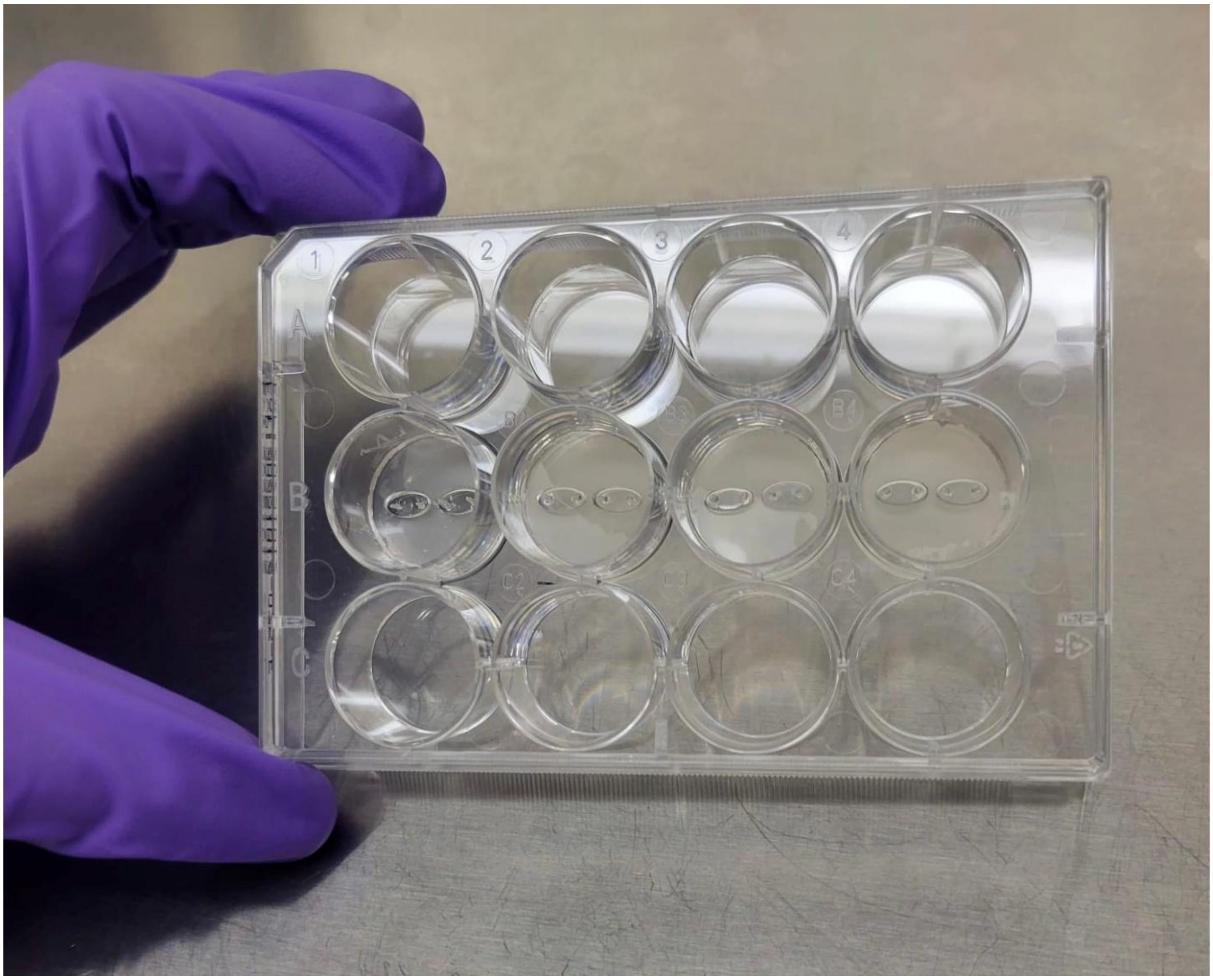


**Fig. S8.** The HOC device fits standard 12-well plates, enabling medium- to high-throughput applications.

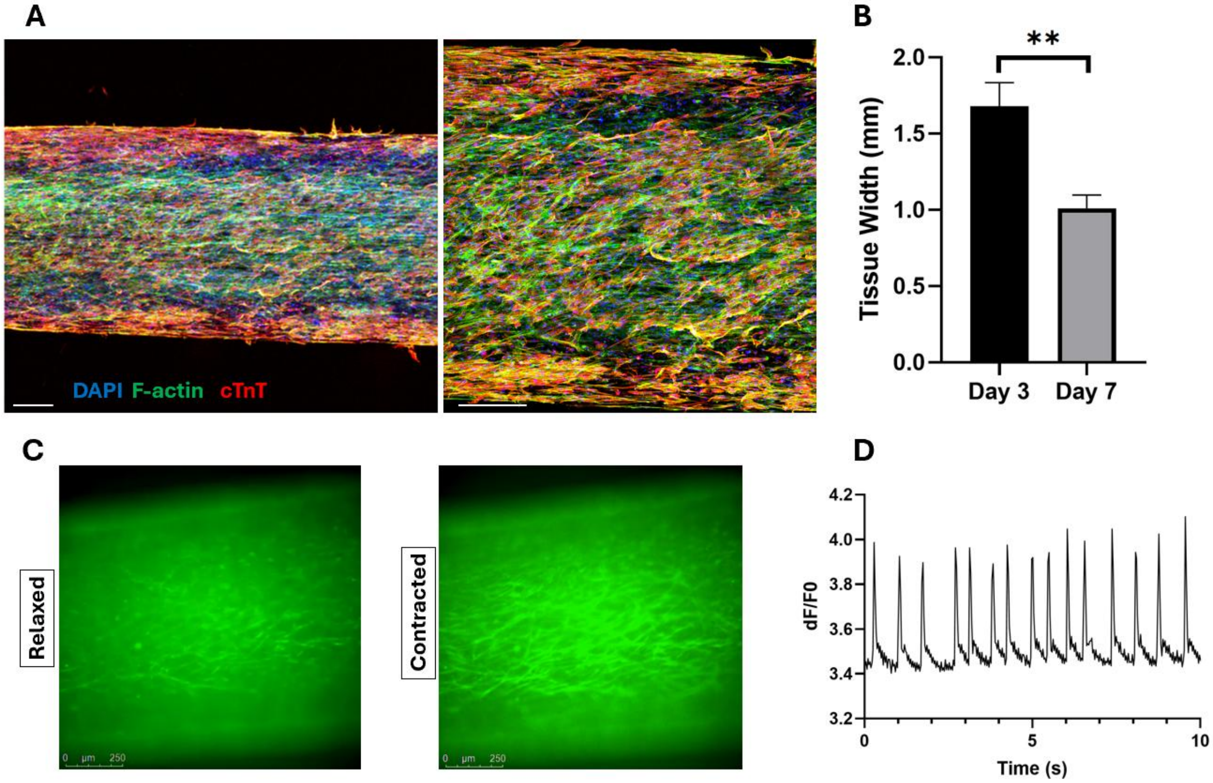


**Fig. S9.** Successful formation of rat EHTs. (A) Elongated cardiomyocytes with organized troponin networks (red); Nuclei and F-actin are shown in blue and red, respectively. Scale bar = 200 µm. (B) Progressive tissue compaction from day 3 to day 7. (C) Functional assessment via dynamic calcium transients, quantified by changes in fluorescence intensity over time. Scale bar = 250 µm.

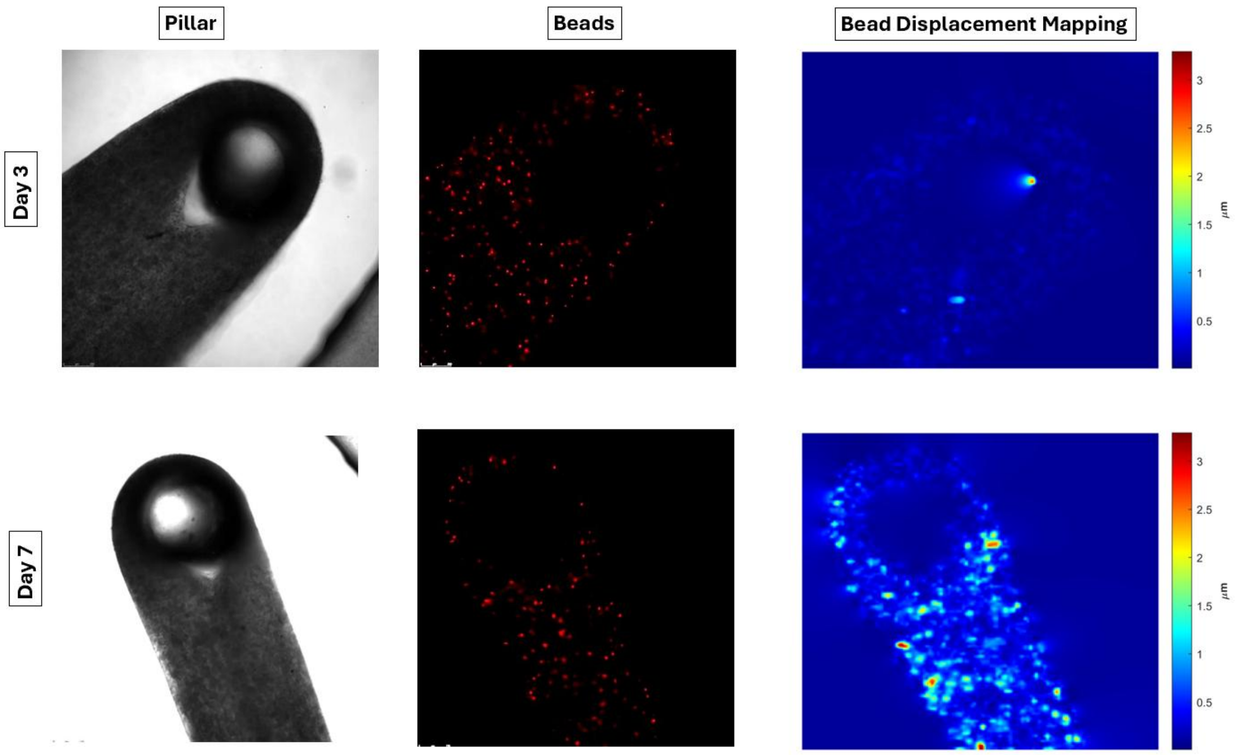


**Fig. S10.** Preliminary validation of real-time contractility mapping using neonatal rat CMs based on tracking bead displacements, showing a progressive increase in contractile strength from day 3 (onset of spontaneous beating) to day 7 (peak of spontaneous beating).

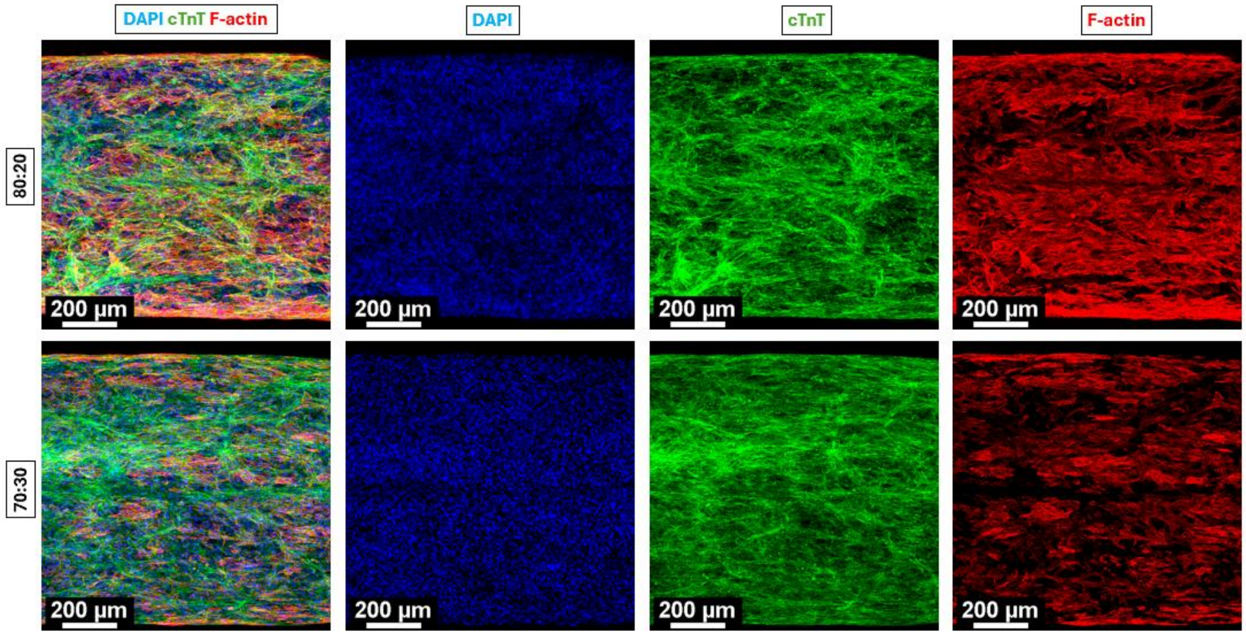


**Fig. S11.** Confocal images of the IF stained EHTs with different ratios of CMs:CFs (80:20 and 70:30), acquired using 10x objective. The images show cTnT in green, DAPI in blue, and F-actin in red.

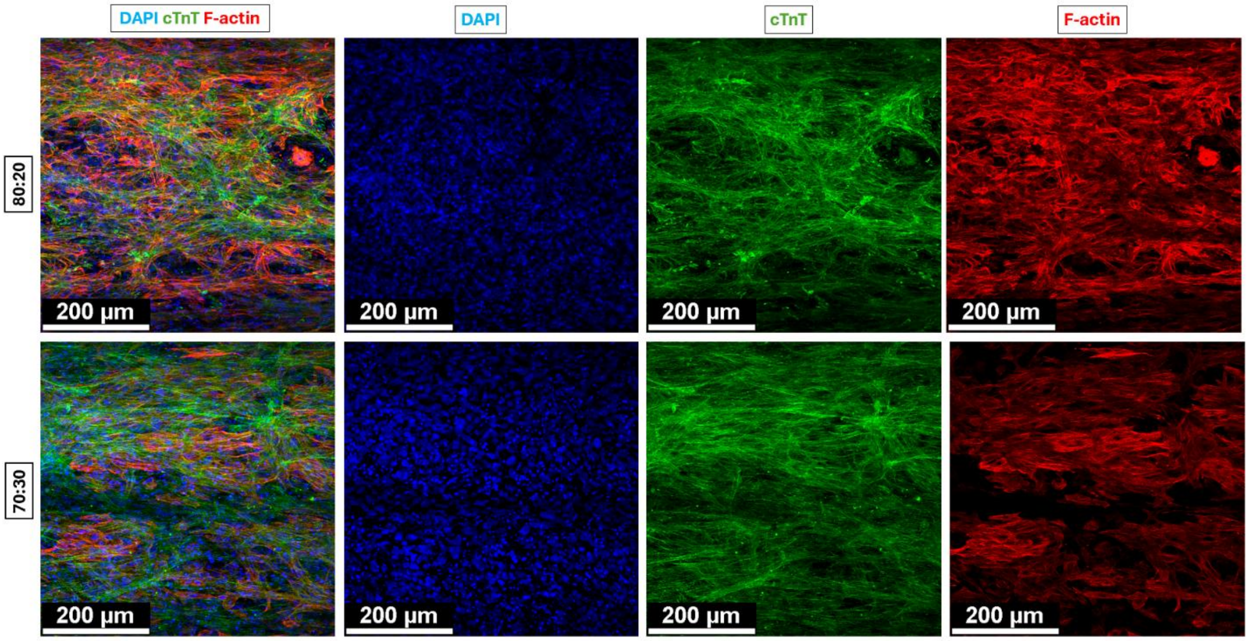


**Fig. S12.** Confocal images of the IF stained EHTs with different ratios of CMs:CFs (80:20 and 70:30), acquired using 20x objective. The images show cTnT in green, DAPI in blue, and F-actin in red.

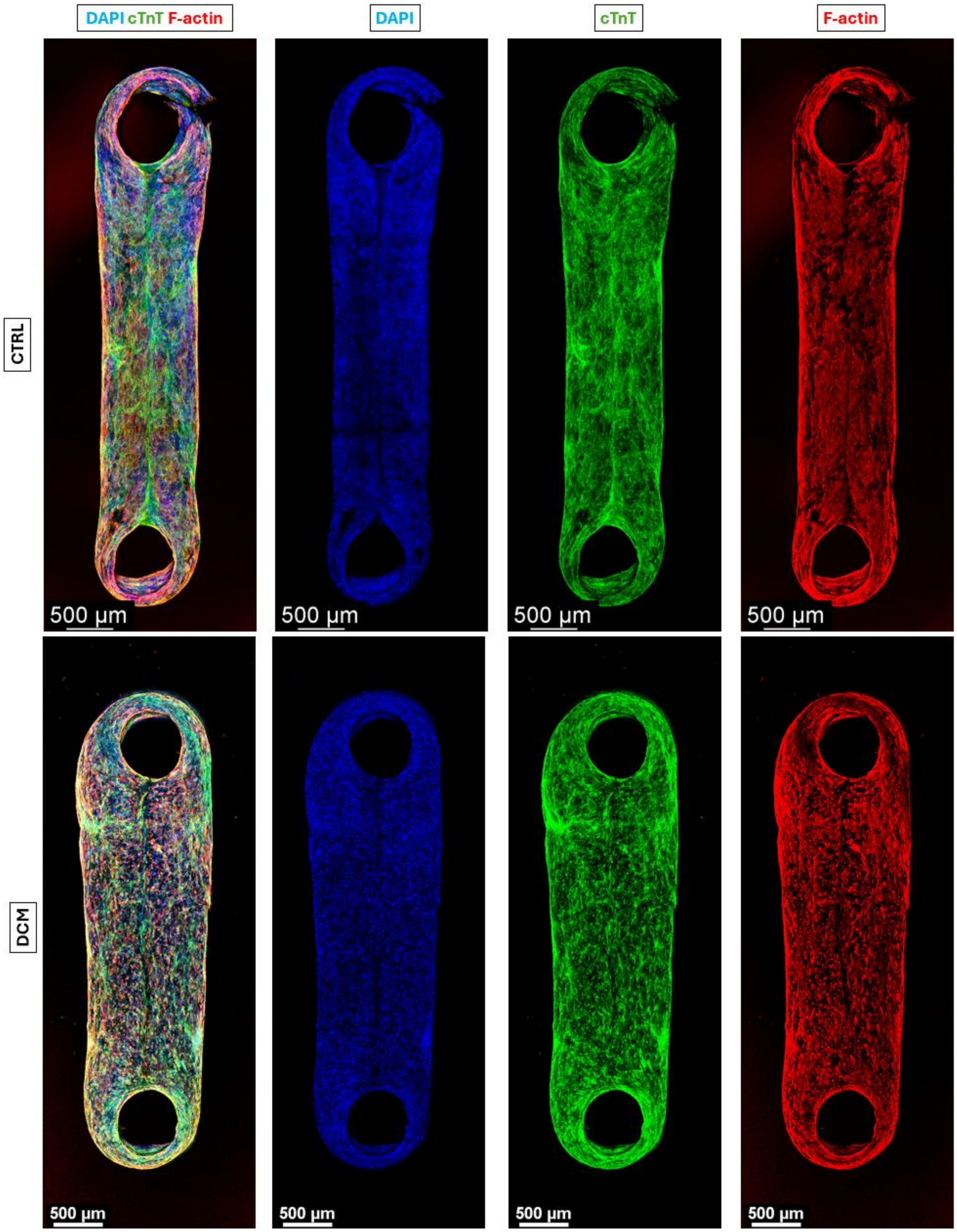


**Fig. S13.** Confocal images of the whole-mount stained control and DCM tissues, showing cTnT in green, DAPI in blue, and F-actin in red.

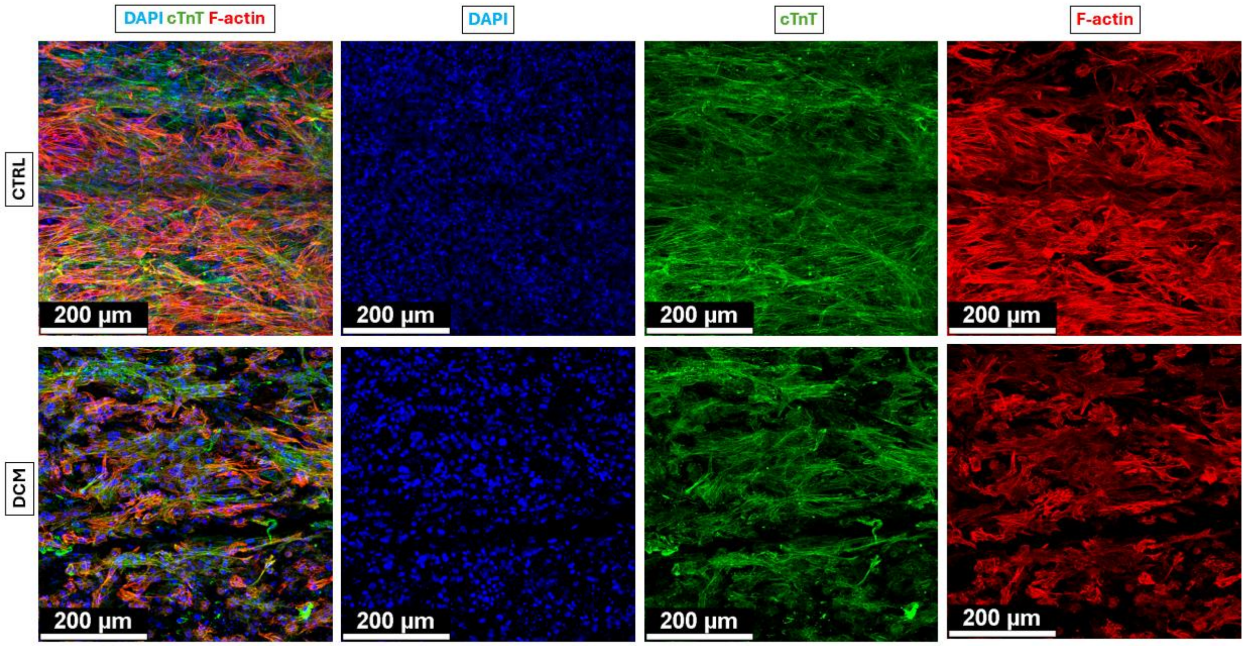


**Fig. S14.** Confocal images of the IF stained control and DCM tissues, acquired using 20x objective. The images show cTnT in green, DAPI in blue, and F-actin in red.

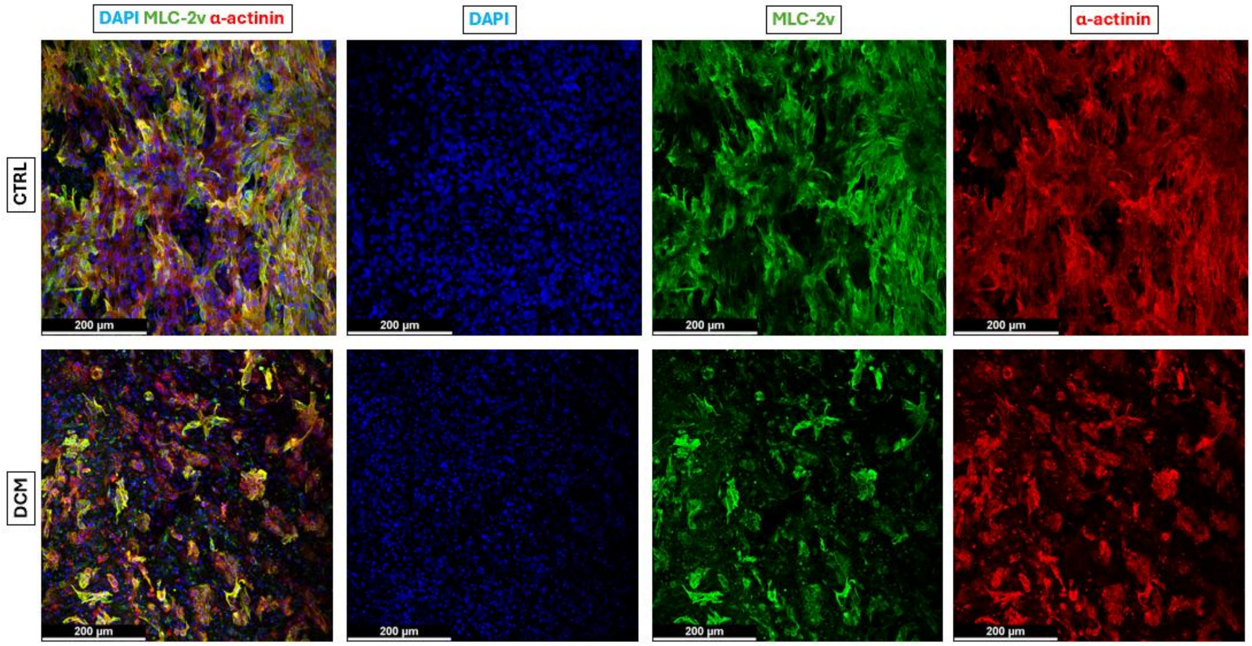


**Fig. S15.** Confocal images of the IF stained control and DCM tissues, acquired using 20x objective. The images show MLC-2v in green, DAPI in blue, and α-actinin in red.

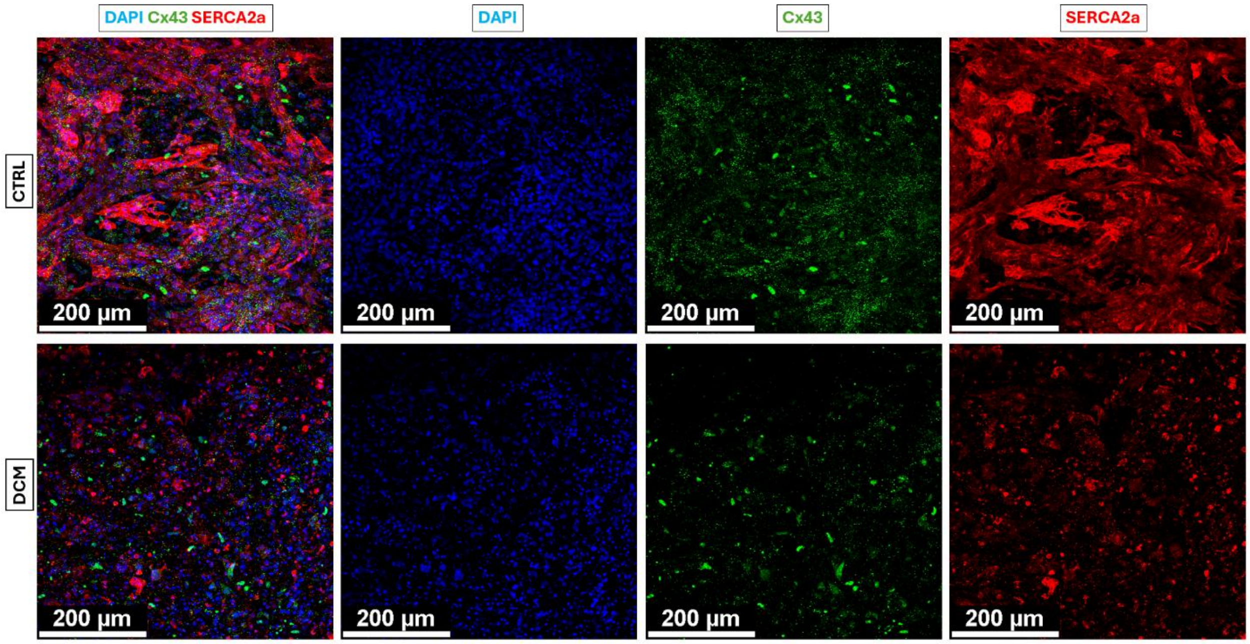


**Fig. S16.** Confocal images of the IF stained control and DCM tissues, acquired using 20x objective. The images show Cx43 in green, DAPI in blue, and SERCA2a in red.

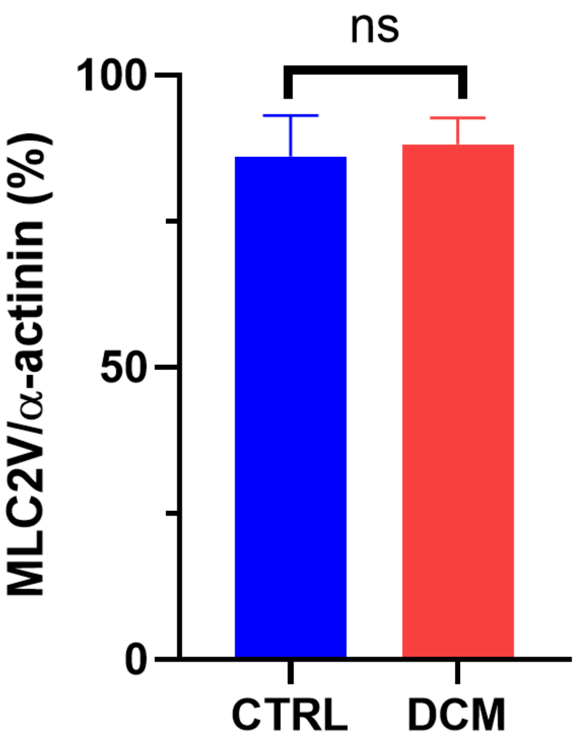


**Fig. S17.** Ratio of MLC-2v to α-actinin density in control and DCM tissues.

**Supplementary Movies**

**Movie S1:** Spontaneous beating of control hiPSC-CM monolayers on day 18 post-differentiation.

**Movie S2:** Spontaneous beating of DCM hiPSC-CM monolayers on day 18 post-differentiation.

**Movie S3:** Representative video of bead movements in EHTs with an 80:20 ratio of CMs to CFs.

**Movie S4:** Representative video of bead movements in EHTs with a 70:30 ratio of CMs to CFs.

**Movie S5:** Representative video of tissue contraction in control EHTs before NE treatment.

**Movie S6:** Representative video of tissue contraction in control EHTs after NE treatment.

**Movie S7:** Representative video of tissue contraction in DCM EHTs before NE treatment.

**Movie S8:** Representative video of tissue contraction in DCM EHTs after NE treatment.
